# Facilitation of sensory axon conduction to motoneurons during cortical or sensory evoked primary afferent depolarization (PAD) in humans

**DOI:** 10.1101/2021.04.20.440509

**Authors:** K. Metz, I. Concha-Matos, Y. Li, B. Afsharipour, C.K. Thompson, F. Negro, DJ. Bennett, MA. Gorassini

## Abstract

Sensory and corticospinal (CST) pathways activate spinal GABAergic interneurons with axo-axonic connections onto proprioceptive (Ia) afferents that depolarize these afferents (termed primary afferent depolarization, PAD). In rodents sensory-evoked PAD is produced by GABA_A_ receptors at nodes of Ranvier in Ia-afferents, rather than at presynaptic terminals, and facilitates action potential propagation to motoneurons by preventing branch point failures, rather than causing presynaptic inhibition. Here we examined if PAD likewise facilitates the Ia-afferent mediated H-reflex in humans by evoking PAD with both sensory and CST stimulation. H-reflexes in several lower limb muscles were facilitated by prior conditioning from low-threshold proprioceptive, cutaneous or CST pathways, with a similar time course (∼200 ms) to the PAD measured in rodent Ia-afferents. Long trains of repeated cutaneous or proprioceptive afferent stimulation produced long-lasting facilitation of the H-reflex for up to 2 minutes, consistent with the tonic depolarization of rodent Ia-afferents mediated by nodal 5-GABA receptors for similar stimulation trains. Facilitation of the conditioned H-reflexes was not mediated by direct facilitation of the motoneurons because isolated stimulation of sensory or CST pathways did not modulate the firing rate of tonically activated motor units in tested muscles. Furthermore, cutaneous conditioning increased the firing probability of a single motor unit during the H-reflex without increasing its firing rate at this time, indicating that the underlying excitatory postsynaptic potential (EPSP) was more probable, but not larger. These results are consistent with sensory and CST pathways activating nodal GABA_A_ receptors that reduce intermittent failure of action potentials propagating into Ia-afferent branches.

**Key Points Summary:**

- The control of posture and movement requires peripheral sensory feedback, which was previously thought to be inhibited by specialized GABAergic neurons in the spinal cord.
- Based on new findings in rodents, we provide evidence in humans that sensory and corticospinal pathways that likely activate these GABAergic pathways facilitate, rather than inhibit, the flow of sensory feedback in afferents that carry information about body position, movement and effort.
- These new findings of how sensory and descending pathways facilitate this sensory feedback to spinal motor neurons can now be applied to people with injury to the brain or spinal cord where these GABA neurons are affected, allowing us to understand how altered sensory control may affect residual motor function and the production of involuntary muscle spasticity.

## Introduction

Peripheral sensory pathways in the spinal cord regulate the transmission of action potentials in other sensory axons through a network of interneurons that release gamma-aminobutyric-acid (GABA) onto these axons. Specifically proprioceptive, cutaneous or pain afferents activate excitatory glutamatergic interneurons, which in turn synapse onto specialized GABAergic interneurons (termed GABA_axo_ neurons; GAD2+) with axo-axonic contacts onto other afferents, forming the classic tri-synaptic pathway described in (Jankowska *et al*., 1981; Alvarez, 1998; Lalonde & Bui, 2021). The activation of GABA_A_ receptors on these sensory axons produces a local depolarization of the afferent due to an outward flow of chloride ions (Gallagher *et al*., 1978) [also reviewed in (Rudomin & Schmidt, 1999; Willis, 1999)]. Although paradoxically excitatory, this GABA_A_ receptor-mediated depolarization (referred to as primary afferent depolarization, PAD) was previously thought to shunt or inactivate action potentials invading the afferent terminal, thereby inhibiting neurotransmitter release to produce presynaptic inhibition (Willis, 2006). This inhibitory role was postulated because the PAD evoked by a flexor afferent followed a similar time course to the suppression of excitatory postsynaptic potentials (EPSP) in an extensor motoneuron when conditioned by the same flexor afferent (Frank & Fortes, 1957; Eccles *et al*., 1961; Willis, 1999). Because there appeared to be no direct effects of the conditioning flexor nerve stimulation on the motoneuron, a presynaptic inhibitory mechanism of PAD on afferent transmission was assumed and this idea has prevailed over the past 60 years (Willis, 2006; Zimmerman *et al*., 2019). However, recent studies reveal that GABA_A_ receptors are generally not on Ia afferent terminals (Alvarez *et al*., 1996; Lucas-Osma *et al*., 2018; Hari *et al*., 2021) and do not depolarize the Ia afferent terminals during PAD (Lucas-Osma *et al*., 2018; Hari *et al*., 2021), conflicting with the concept of presynaptic inhibition. Instead, GABA_A_ receptors are found mostly at the nodes of Ranvier (nodes) in the many large, myelinated branches of Ia afferents in the spinal cord, especially near branch points throughout the dorsal and ventral spinal cord. Accordingly, a novel role of GABA in facilitating, rather than inhibiting, afferent conduction has been proposed where activation of GABA_A_ receptors in the afferent nodes produces a local depolarization that facilitates adjacent sodium channels to secure action potential propagation and decreases downstream branch point failure (termed nodal facilitation) (Hari *et al*., 2021). In the dorsal horn of rodents, these GABA_A_ receptors on the Ia afferent nodes are readily activated by cutaneous or pain afferent pathways and produce nodal facilitation, rather than presynaptic inhibition, of the Ia-mediated monosynaptic reflex (Hari *et al*., 2021). Our preliminary data from the soleus muscle suggests that this may also occur in humans following cutaneous afferent conditioning to evoke PAD (Hari *et al*., 2021). In this paper we expand these studies to provide further evidence for nodal facilitation in human Ia afferents by examining additional muscle groups with various forms of conditioning stimuli to induce PAD. The conditioning included stimulation of proprioceptive and cutaneous afferents and descending corticospinal (CST) motor pathways, the latter which directly synapses onto GABA_axo_ neurons (Ueno *et al*., 2018) and thus, should produce a PAD as shown for afferents in the dorsal horn following trains of motor cortex stimulation (Carpenter *et al*., 1963).

Previous studies in humans have suggested that cutaneous and CST pathways reduce inhibition in Ia afferents produced by a conditioning proprioceptive stimulation, and this was argued to be mediated by dis-facilitating the GABA_axo_ interneurons within the classic tri-synaptic PAD pathway (Rudomin *et al*., 1983), effectively causing a removal of presynaptic inhibition (Berardelli *et al*., 1987; Iles & Roberts, 1987; Nakashima *et al*., 1990; Iles, 1996; Meunier & Pierrot-Deseilligny, 1998; Aimonetti *et al*., 2000). Specifically, as in animal experiments, conditioning by a proprioceptive antagonist nerve was used to suppress the agonist H-reflex and this was assumed (likely incorrectly) to be a demonstration of presynaptic inhibition of the Ia afferents mediating the H-reflex. Then, a prior activation of cutaneous or CST pathways before the conditioning antagonist nerve stimulation was found to reduce this H-reflex suppression. It was proposed that these cutaneous and CST pathways reduced PAD and presynaptic inhibition of the Ia afferents (H-reflexes) by decreasing activity in the GABA_axo_ interneurons. However, recent direct recordings from Ia afferents indicate that a cutaneous conditioning stimulation increases, rather than decreases, GABA_axo_ activity and PAD (Lucas-Osma *et al*., 2018; Hari *et al*., 2021). Thus, we propose a simpler mechanism whereby cutaneous and CST pathways *facilitate* conduction in Ia afferents by activating the GABA_axo_ interneurons that mediate nodal depolarizations to reduce action potential failure at dorsally located branch points. This would not only explain the older data that show cutaneous and CST conditioning causes a reduced inhibition of the H-reflex, but also account for the direct facilitation of the Ia afferents mediating the H-reflex by these conditioning pathways. Here we provide further evidence to support this view.

There are two types of PAD that have been measured in proprioceptive and cutaneous afferents (Rudomin & Schmidt, 1999; Willis, 1999; Delgado-Lezama *et al*., 2013; Lucas-Osma *et al*., 2018; Hari *et al*., 2021). The first is a short-duration (phasic) depolarization lasting for 100 to 200 ms that is evoked from a brief stimulation train (1-3 pulses at 200 Hz) of low threshold proprioceptive, cutaneous or pain afferents (Eccles *et al*., 1962a; Willis, 2006; Lucas-Osma *et al*., 2018). This phasic PAD is mediated by synaptic GABA_A_ receptors with α1, α2 and γ2 subunits that are located adjacent to sodium channels in afferent nodes (Hari *et al*., 2021). The second type of PAD is a longer-duration (tonic) depolarization that lasts for 10’s of seconds (Eccles *et al*., 1962b) and is activated by longer trains of stimulation, most effectively from a fast stimulation train (0.5 s at 200 Hz) or from a relatively slower but longer frequency train (20 s at 0.2 to 2 Hz) (Lucas-Osma *et al*., 2018).

This tonic PAD is mediated by the activation of extra-synaptic GABA receptors with 5α subunits (5 GABA receptors) that are also located near nodal sodium channels and is α specifically reduced by the 5 GABA receptor antagonist L655708 (Lucas-Osma *et al*., 2018; Hari *et al*., 2021).

Importantly, the time course of phasic PAD (100-200 ms), and the reduction in branch point failure it produces in the Ia afferent, is reflected in the time course of motoneuron EPSP potentiation mediated by the facilitated Ia afferents (Hari *et al*., 2021). Thus, in the present study we first examined in human participants if the facilitation of Ia-mediated H-reflexes had a similar time course to the phasic PAD measured in rodent Ia afferents. To do this we measured the facilitation of the H-reflex (as a measure of Ia afferent conduction) produced by a brief peripheral sensory or CST conditioning stimulation at interstimulus intervals (ISIs) between 0 to 200 ms. Second, to determine if we could induce long-lasting increases in H-reflexes indicative of tonic PAD in the Ia afferents, we used long trains of cutaneous stimulation (0.5 to 10 s) to produce tonic PAD, which is thought to result from GABA spillover in the dorsal horn and subsequent long-lasting activation of extra-synaptic 5 GABA receptors and tonic PAD (Lucas-Osma *et al*., 2018). Since Lucas-Osma *et al*., 2018 found that faster cutaneous stimulation trains were more effective in inducing tonic PAD, we examined if fast compared to slower trains produced more long-lasting H-reflex facilitation.

In all experiments, special attention was given to ensure there were no direct postsynaptic effects on the motoneuron produced by the sensory or CST conditioning stimulation to ensure that any facilitation of H-reflexes was likely presynaptic in origin. To rule out direct effects on the motoneuron pool, we examined if the conditioning stimulation itself modulated the firing rate of tonically firing motor units in the test muscle, because the generation of action potentials in a motoneuron is very sensitive to small changes in synaptic inputs, even at distal dendrites, making firing rate a sensitive measure of postsynaptic effects on the motoneuron (Powers & Binder, 2001).

Lastly, we measured changes in the firing rate and discharge probability of single motor units (motoneurons) activated during the H-reflex to estimate the underlying Ia-EPSP by using the peristimulus frequencygram (PSF) method (Turker & Powers, 2005). This allowed us to examine whether spontaneous failures in branch point conduction and EPSPs decreased with PAD. Remarkably, with the PSF method we found that increases in H-reflex amplitude from cutaneous conditioning (and PAD) were not associated with an increase in the underlying Ia-EPSP size, ruling out changes in presynaptic neurotransmitter release and/or postsynaptic effects that should produce *graded* changes in the size of the EPSP. Instead, the all-or-none probability of motor unit discharge during the EPSP increased, suggesting that the underlying EPSP was more probable, but not larger. This is consistent with results from the rat where cutaneous-evoked PAD increases the probability of Ia afferent conduction through branch points and resultant unitary EPSPs (Hari *et al*., 2021). Taken together, these experiments provide evidence for nodal facilitation in the Ia afferent by sensory and CST pathways which helps to overcome intermittent branchpoint failure.

## Methods

### Ethical Approval

Experiments were approved from the Human Research Ethics Board at the University of Alberta (Pro 00078057) and performed with informed consent of the participants. Our sample comprised of 35 participants (18 male) with no known neurological injury or disease, ranging in age from 18 to 57 years (27.0 ± 9.5, mean ± standard deviation).

### Experimental Set up

Participants were seated in a reclined, supine position on a padded table. The right leg was bent slightly to access the popliteal fossa and padded supports were added to facilitate complete relaxation of all leg muscles because CST activation could potentially activate spinal GABA circuits (Ueno *et al*., 2018). For the transcranial magnetic stimulation (TMS) experiments, participants sat in a padded chair with the right leg slightly extended to 100° at both the knee and ankle joint and the foot was strapped to a supporting platform. The upper leg was also supported with straps and padding as above. The head was supported by a headrest to allow minimal movement during TMS. During H-reflex recordings, participants were asked to rest completely with no talking, hand or arms movements.

### Surface EMG recordings

To measure M-wave and H-reflexes, a pair of Ag-AgCl electrodes (Kendall; Chicopee, MA, USA, 3.2 cm by 2.2 cm) was used to record surface EMG from the soleus, tibialis anterior (TA), abductor hallucis (AbHal), biceps femoris, medial gastrocnemius and vastus lateralis (Quad) muscles with a ground electrode placed just below the knee. The EMG signals were amplified by 200 to 1000 and band-pass filtered from 10 to 1000 Hz (Octopus, Bortec Technologies; Calgary, AB, Canada) and digitized at 5000 Hz using Axoscope 10 hardware and software (Digidata 1400 Series, Axon Instruments, Union City, CA). To examine if the conditioning inputs had any direct effects on the motoneuron pool, the surface electrodes were also used to record single motor unit activity in the soleus and AbHal muscles by placing them on the border of the muscle as per (Matthews, 1996).

Motor unit activity from the soleus muscle was also recorded at higher levels of contraction using a high-density surface EMG electrode (OT Bioelettronica, Torino, Italy, semi-disposable adhesive matrix, 64 electrodes, 5×13, 8 mm inter-electrode distance) with differential and ground electrodes wrapped above the ankle and below the knee respectively. Signals were amplified (150 times), filtered (10 to 900 Hz) and digitized (16 bit at 5120 Hz) using the Quattrocento Bioelectrical signal amplifier and OTBioLab+ v.1.2.3.0 software (OT Bioelettronica, Torino, Italy). The EMG signal was decomposed into single motor units using a convolutive blind source separation algorithm implemented in MatLab R2020b with additional quality assessment and accuracy improvement of the automatically decomposed motor unit pulse trains as described previously (Negro *et al*., 2016; Martinez-Valdes *et al*., 2017; Afsharipour *et al*., 2020). Single motor units that were measured from the surface EMG were discriminated visually.

### Nerve Stimulation to evoke homonymous and heteronymous H-reflexes

The tibial nerve (TN) was stimulated using a constant current stimulator (1 ms rectangular pulse width, Digitimer DS7A, Hertfordshire, UK) to evoke a homonymous H-reflex in the soleus and AbHal muscles. After searching for the TN with a surface probe, an Ag-AgCl cathode electrode (Kendall; Chicopee, MA, USA, 2.2 cm by 2.2 cm) was placed in the popliteal fossa, with the anode electrode (Axelgaard; Fallbrook, CA, USA, 5 cm by 10 cm) placed on the patella. If an AbHal H-reflex was not readily evoked from TN stimulation behind the knee, the posterior TN was stimulated below the medial malleolus. A heteronymous H-reflex (> 100 μV) was evoked in the biceps femoris muscle by stimulating the nerve to medial gastrocnemius (MG) in the popliteal fossa (1 ms pulse width, ∼3.3 x M-wave threshold measured in MG muscle). A homonymous H-reflex was evoked in the vastus lateralis (Quad) muscle by stimulating the femoral nerve in the femoral triangle, also with a 1 ms pulse width. Stimulation intensity for the homonymous H-reflexes was set to evoke a test (unconditioned) H-reflex below half maximum on the ascending phase of H-reflex recruitment curve (∼30% of the maximum H-reflex) to reduce the possibility of evoking polysynaptic reflexes and observing increases in Ia conduction (Hari *et al*., 2021). H-reflexes were evoked every 5 seconds to minimize post-activation depression of the Ia afferents (Hultborn *et al*., 1996). At least 20 test H-reflexes were evoked before conditioning to establish a steady baseline since activation of the Ia afferent itself could also activate spinal GABA networks and facilitate H-reflexes (i.e., self-priming, Hari *et al*., 2021). All H-reflexes were recorded at rest except during the firing probability experiments described below.

### Sensory and CST conditioning of H-reflexes to produce phasic PAD

*Cutaneous and proprioceptive conditioning stimulation:* To condition the H-reflex by mainly cutaneous afferents, the medial (cutaneous) branch of the deep peroneal nerve (cDPN) was stimulated on the dorsal surface of the ankle using a bipolar arrangement (Ag-AgCl electrodes, Kendall; Chicopee, MA, USA, 2.2 cm by 2.2 cm). A short train (3 pulses, 200 Hz for 10 ms) of cDPN stimulation was applied at intensities corresponding to perception threshold but below radiating threshold (3.0 to 7.4 mA) to avoid direct activation of motoneurons as assessed below. A train of pulses was used for the sensory nerve stimulation because it evokes a larger phasic PAD in Ia afferents compared to single pulse stimulation (Eccles *et al*., 1962c). Approximately 20 baseline SOL H-reflexes were elicited to ensure the H-reflex was stable, followed by 7 conditioned H-reflexes at one of the ISIs (0, 30, 60, 80, 100, 150 or 200 ms). Following this, 7 unconditioned H-reflexes were evoked to re-establish baseline and another run of 7 conditioned H-reflexes was applied at another randomly chosen ISI. This was repeated until all ISIs were applied. To condition the H-reflex by mainly low-threshold proprioceptive afferents, the common peroneal nerve (CPN) was stimulated in a bipolar arrangement just below the head of the fibula (3 pulses at 200 Hz) at 1.0 x motor threshold measured in the tibialis anterior muscle, being careful to elicit a pure dorsiflexion response.

*CST conditioning stimulation:* The CST to the soleus, AbHal or Quad motoneuron pool was activated by applying TMS to the contralateral motor cortex using a custom made figure-of-eight batwing coil [P/N 15857; 90 mm diameter, (Nielsen & Petersen, 1994)] that was connected to a Magstim 200 stimulator (Magstim; Dyfed, UK). The coil was typically positioned 2 cm lateral to vertex to target the soleus and AbHal muscles. Active motor threshold (AMT) was determined by the lowest-intensity TMS pulse that produced a discernable and reproducible motor evoked potential (MEP) in the tested muscle while the participant held a small voluntary contraction. The TMS intensity was set to 0.9 x AMT in the resting muscle to avoid direct activation in the motoneuron. H-reflexes were conditioned by TMS at the same ISIs as for the short-duration sensory conditioning experiments, in addition to the 250 and 300 ms ISIs given the longer duration of facilitation in some participants.

*Data analysis:* For both the sensory and CST conditioning experiments, the unrectified, peak-peak amplitude of the 7 test H-reflexes immediately preceding the 7 conditioned H-reflexes for a given ISI were averaged together because test H-reflexes could grow over time (self-priming, Hari *et al*., 2021). The effect of the conditioning stimulation on the test H-reflex was measured using the formula: % change H-reflex = ([(conditioned H - test H)/test H] *100%). Data were also analyzed by averaging the amplitude of all test H-reflexes in a trial and this provided similar results throughout so only calculations of % change using the immediately preceding test H-reflexes are reported here. The mean % change H-reflex for each ISI was averaged across participants. Because the profile of H-reflex facilitation could be variable between participants, the maximum or peak % change H-reflex, irrespective of ISI, was also averaged across participants.

*Postsynaptic effects of conditioning stimuli*: To determine if there were any direct effects on the motoneurons from the conditioning stimulation at the time the H-reflexes were evoked, we measured if the sensory or CST stimulation produced any changes in the tonic firing rate of single motor units or changes in the amplitude of the rectified surface EMG. Single motor units were activated in the soleus or AbHal muscle while the participant held a small voluntary contraction around 5% of maximum. Both auditory and visual feedback was used to keep the firing rates of the units steady while the conditioning cutaneous or CST stimulation was applied every 3 to 5 seconds. The firing rate profiles from many stimulation trials were superimposed and time-locked to the onset of the conditioning stimulation to produce a peri-stimulus frequencygram (PSF) as done previously (Turker & Powers, 2005; Norton *et al*., 2008). A mean PSF profile was produced by averaging the frequency values into 20 ms bins. The mean rate in each 20 ms bin was compared to the mean firing rate measured in a 100 ms window before the conditioning stimulation and expressed as: % change PSF = ([(mean bin rate - mean pre-stimulus rate)/mean pre-stimulus rate] *100%). A similar binning process and averaging was done for the rectified surface EMG values using a 100 ms window before the conditioning stimulation to measure the pre-stimulation background EMG.

*Motoneuron and Ia afferent excitability during conditioned H-reflex:* We wanted to estimate the excitability of the motoneuron (as reflected in the PSF and rectified EMG), and PAD in the Ia afferent, at the time the conditioned H-reflexes were activated at the spinal cord. First, to align motoneuron excitability, the conditioned H-reflex values, as a function of the various ISIs (see bottom graph, Fig. 1), were shifted to the right of the onset of the conditioning stimulation in the PSF (light blue trace) and EMG profiles (not shown) by an amount equivalent to the latency of the H-reflex (∼30 ms, length of thin black arrow in Fig. 1). The rightward shift accounted for the time it takes the Ia afferent volley to reach the spinal cord plus the time it takes the motoneuron response to reach the muscle where the H-reflex is measured. Thus, the PSF (or EMG) values occurring near the shifted conditioned H-reflexes can be used as an estimate of motoneuron excitability at the time the conditioned H-reflexes were activated at the spinal cord. Second, plotting the estimated time course of PAD in a Ia afferent after a conditioning stimulation helps to predict when the conditioned H-reflexes should be facilitated by PAD. Because the conditioning-evoked PAD in the Ia afferent is activated at the cord with a similar delay as the Ia EPSP mediating the H-reflex (top two traces in Fig. 1), a 0 ms ISI corresponds to just before the conditioning afferent stimulation can influence the H-reflex. In contrast, at the 80 ms ISI while PAD is activated in the Ia afferent, the H-reflex can be facilitated (Fig 1; compare pink and blue dots representing H-reflexes of inset at 0 and 80 ms ISI respectively). Moreover, if the mean firing rate of the motor units (PSF, light blue line) near the 80 ms ISI is at or below the pre-conditioned level (below red line within red double arrow), this would indicate that the facilitation of the conditioned H-reflex at this ISI was *not* mediated by a depolarization of the motoneurons from the conditioning stimulation.

**Figure 1.**
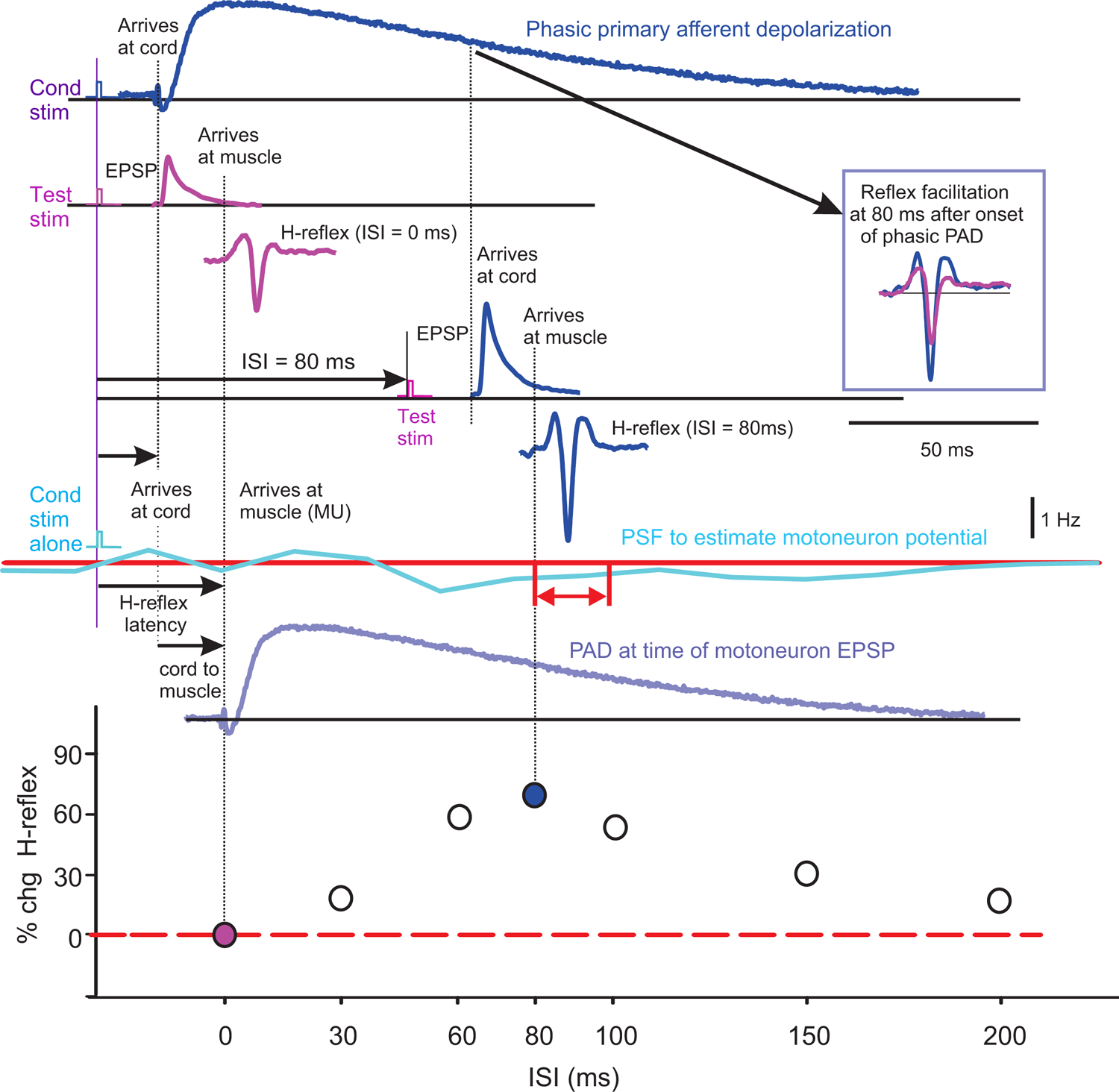
Time course of predicted PAD and its relationship to H-reflex facilitation. Top three traces: *Trace 1*) Example of phasic primary afferent depolarization (PAD) recorded in rat Ia afferent (top dark blue trace). *Trace 2*) Theoretical EPSP recorded in spinal motoneuron and H-reflex recorded in muscle (pink traces) from simultaneous conditioning (Cond) and TN stimulation to produce a conditioned H-reflex as measured for the 0 ms ISI (bottom trace). Time of stimulation marked by rectangular pulses on left. PAD profile taken from intracellular recording in rat Ia afferent (sacral S3) in response to stimulation of an adjacent dorsal root (S4: 1.1 x threshold, 0.1 ms pulse, Hari et al., 2021). *Trace 3*) EPSP (upper) and H-reflex (lower dark blue traces) in response to TN stimulation applied 80 ms after the conditioning stimulation (ISI of 80 ms). **Bottom three traces**: *Trace 4*) Example mean PSF (light blue trace) to represent profile of motoneuron excitability in response to conditioning stimulation alone. Red trace represents mean firing rate before conditioning stimulation was applied. Double red arrow marks the excitability of the motoneuron (potential estimated from the PSF) during activation of the H-reflex. *Trace 5*) Violet trace is profile of PAD shifted to the right by the conduction time from the spinal cord to the muscle to visually line up the excitability of the Ia afferents at the time the motoneuron is being activated during the H-reflex and how this affects the resulting facilitation of the H-reflex recorded at the muscle. *Trace 6*) Percent change in H-reflex amplitude (conditioned H – test H)/test H x 100% at the various ISIs, highlighting the 0 and 80 ms ISIs.

### Cutaneous and proprioceptive facilitation of H-reflexes during tonic PAD

*a) Slow (0.2 Hz) vs moderate (2 Hz) cutaneous frequency trains:* A slow (0.2 Hz) and moderate (2 Hz) frequency train of cDPN stimulation was applied as per animal studies (Lucas-Osma *et al*., 2018). A higher stimulation intensity at 2 times perception threshold (10.6 ± 1.6 mA, n = 16 participants) was used to induce GABA spillover. Following a baseline of 20 test H-reflexes (delivered every 5 s), the slow or moderate cDPN stimulation train was applied along with the test H-reflexes for 10 s, with each cDPN stimulation occurring 500 ms before any H-reflex. Test H-reflexes were evoked for another 120 s to examine aftereffects from the cDPN trains. This stimulation protocol was repeated 3 times for both the 0.2 Hz and 2 Hz stimulation trains in each participant, waiting at least 1 minute at rest between each trial.
*b) Fast (200 Hz) cutaneous frequency train:* A faster (200 Hz) but shorter (500 ms) train of cDPN stimulation was also used to condition the H-reflex as per (Hari *et al*., 2021) using a protocol similar to the slower 10 s trains. Here, the cDPN train was applied 700 ms before the test H-reflex and following this, H-reflexes continued to be evoked for another 90 to 120 s. A very low intensity of stimulation below radiating threshold (3.0 to 4.5 mA, n = 15 participants) was used to ensure the fast frequency stimulation was not painful.
*c) Slow (0.2 Hz) proprioceptive frequency train:* Because repetitive activation of Ia afferents also produces tonic PAD (Hari *et al*., 2021), we examined if there was a buildup of test H-reflexes from repetitive stimulation of the TN afferents alone. In rodents, activation of Ia afferent collaterals activate PAD networks that connect back to the same afferent to produce self-facilitation, which is revealed during low-intensity, repetitive stimulation of Ia afferents (1.1 x afferent threshold, 0.1 Hz, ∼ 30% of maximum EPSP). Thus, we measured the amplitude of test H-reflexes activated every 5 seconds (0.2 Hz) at a low intensity of TN stimulation (∼30% of maximum H-reflex) in 19 participants before any conditioning (cutaneous or CST) stimulation was applied.

*Data analysis*: H-reflexes following the cutaneous conditioning train in experiments *a* and *b* (post-train H) were compared to the average amplitude of the 20 baseline H-reflexes (pre-train H) using the % change formula: [(post-train H – pre-train H)/pre-train H] *100%. The resulting % change H-reflex values were plotted against time and divided into 10 s bins (2 H-reflexes per bin). H-reflexes from all 3 trial runs were grouped together (2 H-reflexes per bin x 3 trials = 6 H-reflexes per bin). The average % change H-reflex in each bin was then averaged across all 16 participants. In experiment *c* the amplitude of each of the first 14 test H-reflexes measured at baseline before any conditioning stimulation was applied (set 1 in Experimental Protocol, Fig. 2) was expressed as a percentage of the average of the 7 test H-reflexes following the first conditioning run (in set 2) where the H-reflexes reached a steady state using the formula: (set 1 H_(1→14)_ /avg set 2 H) * 100%. Each 1^st^ to 14^th^ %H-reflex was then averaged across the 19 participants and plotted against stimulation number (and time).

**Figure 2.**
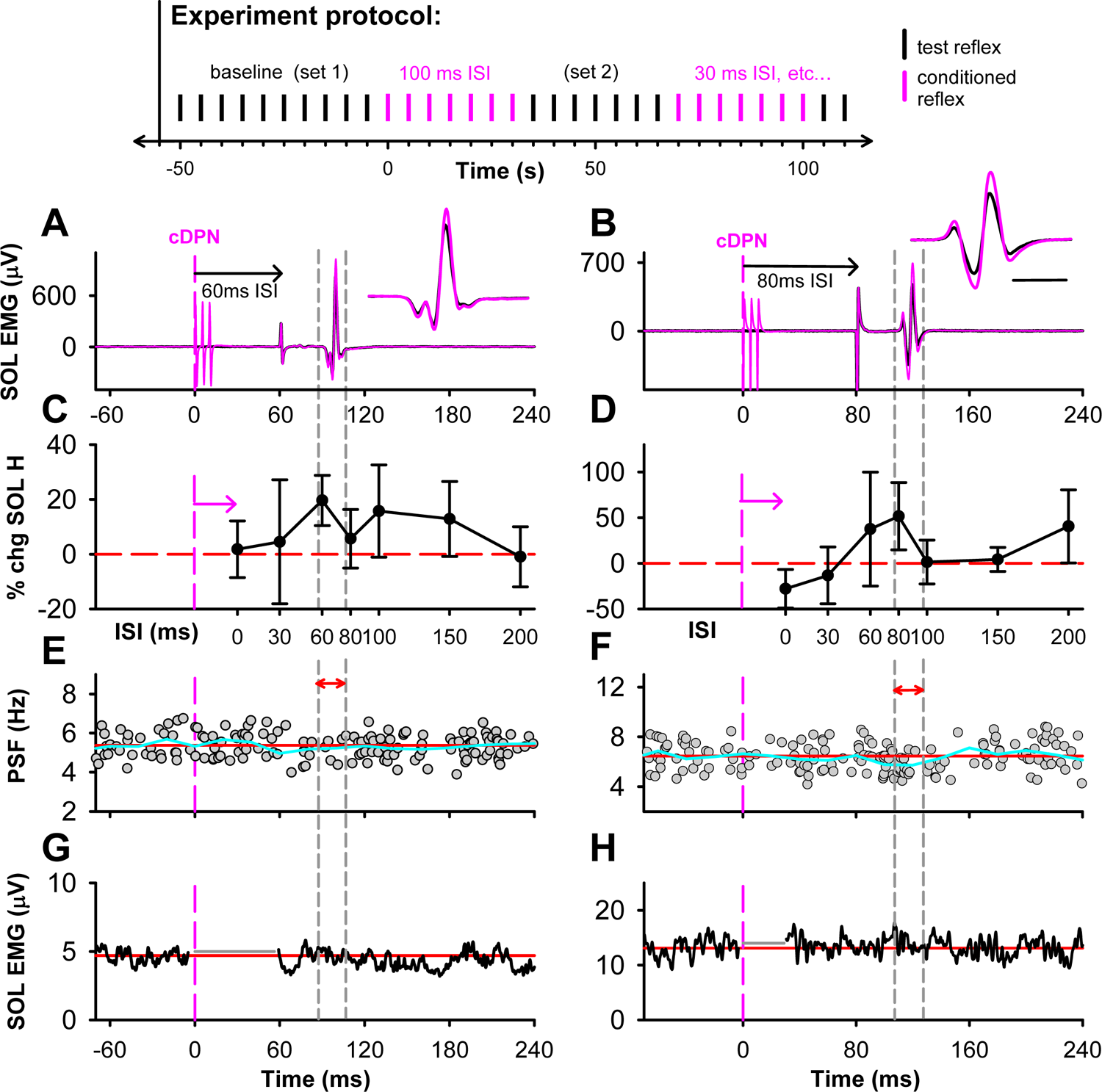
Short-duration H-reflex facilitation by cutaneous inputs. **Top:** Experiment protocol showing alternating sequence of applying test (black) and conditioned (pink) H-reflexes at the various ISIs. **A&B)** Representative soleus (SOL) H-reflex modulation from cDPN conditioning stimulation in 2 representative participants. Average of 7 test (black) and 7 cDPN-conditioned (pink) SOL H-reflexes (200 Hz, 10 ms, 3.5mA and 4.0mA respectively) with TN stimulation at 60 ms in A and 80 ms in B (expanded time scale for H-reflex in insets, time bar = 10 ms). **C&D)** % change of the SOL H-reflex at each ISI (mean ± SD). Peak % change marked by dashed vertical lines. Data points are shifted to the right of the onset of cDPN stimulation (pink dashed vertical line) in E&F. The amount of shift is equal to the onset latency of the H-reflex (length of pink arrow). **E&F**) PSF of a SOL motor unit, time-locked to the time of the cDPN conditioning stimulation alone at 0 ms (vertical pink line) with 110 sweeps in E and 180 sweeps in F. Firing rate averaged into 20 ms bins (blue trace) for comparison to the average pre-stimulus firing rate (red line). **G&H)** Averaged rectified SOL EMG recorded with the cDPN conditioning stimulation alone with 112 sweeps in G and 82 sweeps in H. Stimulation artifact removed (grey horizontal line). Average pre-stimulus EMG marked by the horizontal red line. Note x-axis in A-B, E-H is in time (ms) and C-D in ISI (ms).

### Firing probability of single motor units activated during H-reflex by cutaneous conditioning

The firing probability of single motor units during the H-reflex window (approximately 30 to 40 ms post TN stimulation) was measured with and without cDPN conditioning in 13 participants. In 12 participants, single motor units were identified from high density surface EMG (HDsEMG) and in 1 participant with intramuscular EMG to verify the HDsEMG recordings. The size of the H-reflex was set to just above threshold (5.2±3.9% M_max_) during a small plantarflexion so that single motor units at the time of the H-reflex could be distinguished from the compound H-reflex potential (Yavuz *et al*., 2015; Nielsen *et al*., 2019). In a single trial run, test H-reflexes were evoked every 3-5 s for the first 100 s followed by cDPN-conditioned H-reflexes for the next 100 s using a 200 Hz, 50 ms pulse train (3.0 to 4.5 mA, below radiating threshold) applied 500 ms before each H-reflex.

Approximately 40-50 usable test and conditioned firing rate profiles (PSFs) were produced for a single trial run where the motor units had a steady discharge rate 400 ms before and 600 ms after the cDPN stimulation. Trial runs were repeated 3-6 times to obtain a sufficient number of frequency profiles to construct the PSF (∼200, Norton *et al*., 2008).

*Data analysis*: For each test or conditioned PSF, the probability that a motor unit discharged during the ∼15 ms H-reflex window was measured using the following formula: [(number of discharges during the H-reflex window) / (total number of sweeps) *100%]. The mean background firing rate 100 ms before the TN stimulation with and without conditioning was also measured. For both the test and conditioned trials, the average firing probability during the H-reflex window and the mean background rate were measured for each participant and then averaged across the 13 participants. The mean firing rate during the H-reflex window was also measured as an estimate of EPSP size (Norton *et al*., 2008), and expressed as a % change between the test and conditioned H-reflex trials ([(conditioned rate-test rate)/test rate]*100).

### Statistical Analysis

The % change of various measures (conditioned H-reflex, PSF and rectified EMG) across the different ISIs were compared to a 0% change using either a one-way ANOVA for repeated measures for normally distributed data (determined by the Equal Variance test) or by a Friedman Repeated Measures Analysis of Variance on Ranks for data that was not normally distributed. Post hoc Tukey tests for the ANOVA and Friedman were used to determine which ISIs were significantly different from a 0% change. A two-way ANOVA for repeated measures was used to compare the % change in H-reflexes from the 2 Hz and 0.2 Hz conditioning cutaneous stimulation trains, with frequency and time as factors. Post hoc Tukey tests were used to determine which time bins were significantly different between the two stimulation frequencies. A Mann-Whitney U Test was used to compare group values that were not normally distributed and Student’s t-tests for normally distributed data. Data are presented in figures and in the text as mean ± standard deviation (SD). Significance was set as *p* < 0.05 and n refers to the number of participants tested in each experiment.

## Results

*Cutaneous and proprioceptive facilitation of H-reflexes during phasic PAD:* To explore whether conduction in Ia afferents was facilitated during the relatively long duration of phasic PAD (100 - 200 ms, Hari *et al.,* 2021), we started by examining whether the soleus H-reflex was facilitated by a cutaneous conditioning stimulation (cDPN: 4.1 ± 1.1 mA at perception threshold) at interstimulus intervals (ISI) between 0 and 200 ms (see schematic of experimental protocol, top of Fig. 2). Consistent with the time course of phasic PAD, the soleus H-reflex was facilitated when the brief cDPN stimulation was applied between 60 to 150 ms earlier, as shown for two participants in Figures 2A-C & B-D respectively. The change in the conditioned H-reflex with respect to the test H-reflex is plotted as a function of the ISI (Figs. 2C and D) to evaluate how the facilitation changes with the expected time course of phasic PAD (up to 200 ms; see below). For this analysis, the H-reflex and conditioning cDPN volley are expected to have a similar latency in reaching the spinal cord (see Methods, Fig. 1) and thus, an ISI of 0 ms corresponds to a time just before the expected onset of PAD where appreciable H-reflex facilitation is not expected. Longer ISIs correspond to times during the activation of PAD where Ia and reflex facilitation is expected, as highlighted for the H-reflexes at the 60 and 80 ms ISIs in the two participants (see insets in Figs. 2A-B with corresponding %change values shown between the grey dashed lines in Figs. 2 C-D).

The facilitation of the soleus H-reflex occurred even though the conditioning cDPN stimulation itself (when applied alone) did not facilitate the soleus motoneuron pool as reflected in the mean motor unit firing profile (PSF, light blue line in Figs. 2E and F), remaining close to or slightly below the mean firing rate before the cDPN stimulation was applied (red line), and thus ruling out postsynaptic facilitation of the H-reflex. Likewise, the cDPN stimulation did not produce an increase in the mean rectified EMG, representing the activity of a larger number of motor units and ruling out the recruitment of additional units from the conditioning input (Figs. 2G and H). For comparing the PSF or EMG after the cDPN stimulation (Figs. 2E-H) to the conditioned H-reflex changes (Figs. 2C-D), we shifted the H-reflex-ISI plots by the latency of the H-reflex to determine how the cDPN stimulation affected the excitability of the soleus motoneurons at the time the H-reflex was activated (if at all; see Methods, Fig. 1). As highlighted for the maximally facilitated H-reflexes at the 60 and 80 ms ISIs (between grey dashed lines in Fig. 2), both the PSF and EMG remained close to or slightly below the pre-stimulus values indicating an unfacilitated soleus motoneuron pool when the H-reflexes were evoked and facilitated.

Overall from the 16 participants, the average profile of soleus H-reflex facilitation from the conditioning cDPN stimulation resembled the profile of afferent depolarization (PAD) evoked by a cutaneous afferent stimulation (Hari *et al*., 2021), lasting for ∼150 ms but with a later peak at 80 ms (Figs. 3Ai) and with a significant facilitation of the reflex at the 80 and 100 ms ISIs (see legend for statistics). In contrast, there was no change in the mean PSF or EMG across all the ISIs, again indicating a lack of direct effect from the conditioning cDPN stimulation on the soleus motoneurons. The maximal facilitation of the soleus H-reflex across the different condition-test ISIs in each participant was 42.0 ± 27.4% (*p* < 0.001, Fig. 3B left bar), whereas the average firing rate during the PSF when the maximal H-reflex was evoked at the spinal cord (e.g., within the grey dashed lines in Figs. 2E and F) was reduced slightly compared to the firing rate before the cDPN stimulation (−3.4 ± 4.8%, *p* = 0.020, Fig. 3B middle bar). Although there was no change in the overall rectified EMG (Fig. 3B right bar), in 3 participants the rectified EMG was increased compared to the mean pre-stimulus level but this was likely in response to motor unit synchronization following a preceding EMG suppression, highlighting that caution should be used when using EMG to estimate motoneuron depolarization levels (Turker & Powers, 2005). In summary, the facilitation of the H-reflex by the cDPN stimulation was likely not produced by a direct facilitation of the soleus motoneuron pool at the time of Ia activation based on the PSF and EMG. Rather, facilitation likely occurred at a pre-motoneuron level, potentially due to facilitation of spike propagation on the Ia afferents mediating the H-reflex as demonstrated in the motor unit firing probability experiment detailed below.

**Figure 3.**
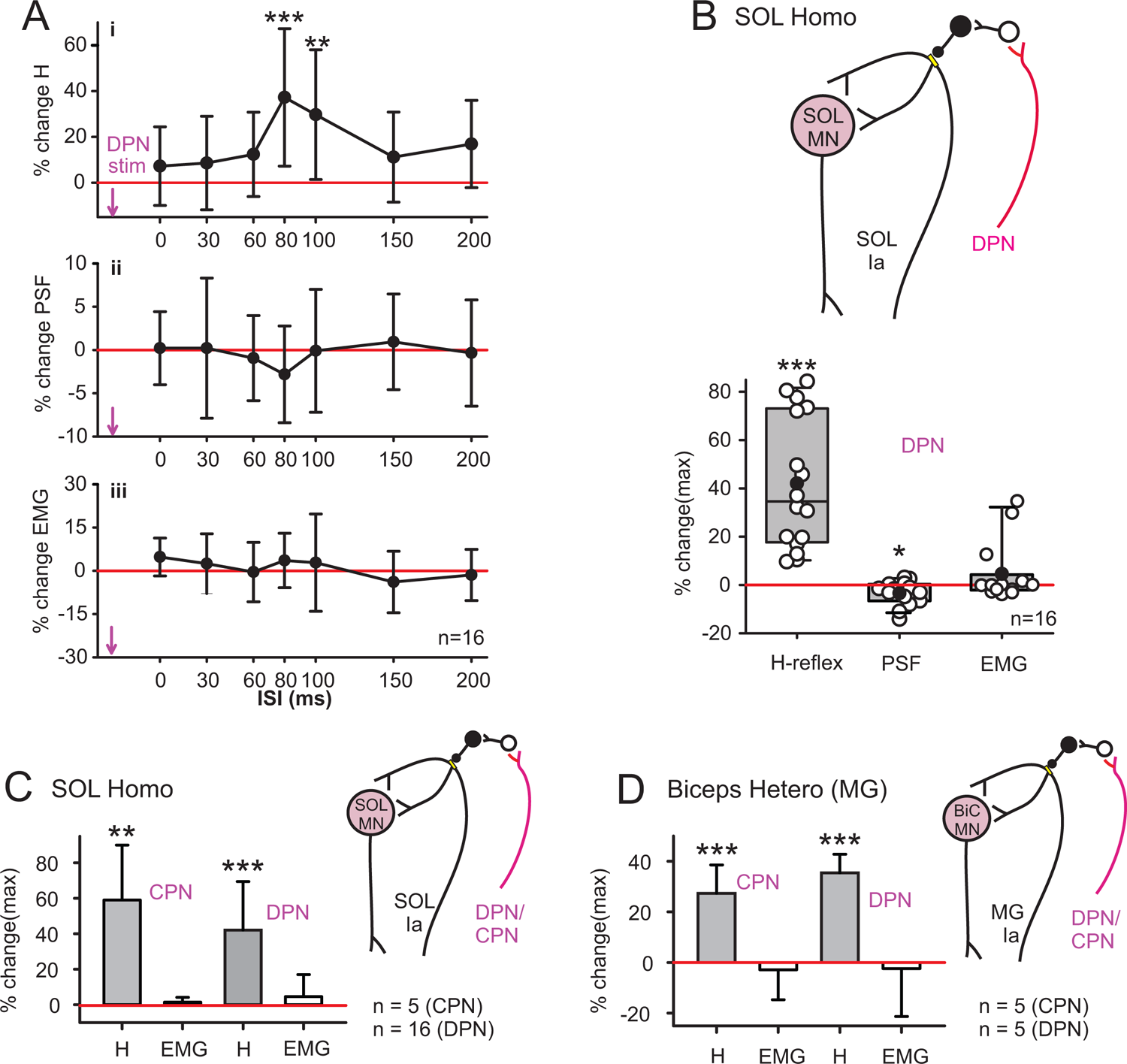
Short-duration H-reflex facilitation by cutaneous inputs: group data. **A**) Average % change in (i) soleus H-reflex, (ii) motor unit firing rate (PSF) and (iii) rectified EMG from a prior cutaneous DPN conditioning stimulation (3 pulses, 200 Hz) at each ISI across the group (n=16). Error bars represent standard deviation about the mean. The average size of the test (unconditioned) SOL H-reflex was 0.86 ± 0.76 mV (11.4 ± 9.2% of Mmax). There was an effect of the conditioning-test ISI on the H-reflex [F(15,7) = 4.3, P < 0.001, one-way ANOVA], being significantly greater than a 0% change at the 80 and 100 ms ISIs (*p* all < 0.01, Tukey) but no effect of ISI for both the mean PSF (Chi-square, DF 7, *p* = 0.52) and EMG [F (13,7) = 1.1, *p* = 0.35], one-way ANOVA). **B)** Left bar: Maximum (peak) % change of the conditioned SOL H-reflex for each participant (white circles), mean represented by black circle and median by horizontal line, 25th and 75th percentiles by the box bounds, and the 95th and 5th percentiles by whiskers. Middle bar: % change PSF at the time bin where the peak conditioned H-reflex would have been activated at the spinal cord (n=16: 5 units recorded using intramuscular or surface EMG, 11 units decomposed from HDsEMG). Right bar: % change of rectified surface EMG as in PSF, no difference from 0 %change (4.6 ± 12.4%, *p* = 0.51). **C)** Maximum % change in soleus H-reflex (H) from conditioning of CPN and DPN, and corresponding % change in EMG. Mean ± SD. **D)** Same as in C but for maximum % change in H-reflex and EMG recorded in biceps femoris muscle. Mann-Whitney U test used for pairwise comparisons to 0 %change with * p < 0.05, ** p < 0.01, *** p<0.001 representing statistically significant difference.

Conditioning with proprioceptive afferents also produced facilitation of both homonymous and heteronymous H-reflexes. A low intensity of stimulation to the common peroneal nerve (CPN, 1.0 x MT), mainly to recruit proprioceptive afferents from the tibialis anterior muscle (antagonist to the soleus), produced a maximum facilitation of soleus H-reflexes of 59.1 ± 30.9% in 5 participants (*p* = 0.008), with no corresponding increase in EMG activity (*p* > 0.50, Fig. 3C). Similar to the cutaneous DPN stimulation (replotted in Fig. 3C), the maximum facilitation of the soleus H-reflex from CPN conditioning occurred between the 60 and 80 ms ISIs (average 68.0 ± 11.0 ms, not shown). The conditioning CPN stimulation also facilitated a heteronymous H-reflex in the biceps femoris muscle (Biceps Hetero) activated by the MG nerve, with a maximum facilitation of 27.4 ± 11.2% (*p* < 0.001, n = 5) that occurred between the 60 and 80 ms ISIs (64.0 ± 8.9 ms, Fig. 3D). Likewise, the cutaneous DPN stimulation facilitated the heteronymous biceps femoris H-reflex by 35.4 ± 7.4% at an average ISI of 60.0 ± 14.1 ms (*p* < 0.001, Fig. 3D), with both low-intensity conditioning stimuli producing no change in the surface EMG reflecting the excitability of the motoneuron pool at the time the H-reflex was evoked (*p* > 0.65, Fig. 3D). Thus, the facilitation of the H-reflex by sensory conditioning not only follows the time course of phasic PAD, but also spatially mimics the widespread distribution of PAD in Ia afferents across many spinal segments from both flexor and extensor muscles (Rudomin & Schmidt, 1999; Willis, 2006).

*CST facilitation of the H-reflex during phasic PAD:* Similar to the action of sensory conditioning, the H-reflex was also facilitated by prior conditioning from the CST activated by TMS to the contralateral motor cortex, which should produce PAD (Carpenter *et al*., 1963). This occurred for the soleus, AbHal and Quad H-reflexes (representative AbHal H-reflex data shown for two participants in Figs. 4A and B, inset). A very weak TMS intensity (0.9 x AMT applied at rest) was used to condition the resting H-reflex to avoid direct facilitation of the motoneuron pool. The profile of AbHal H-reflex facilitation from TMS conditioning at the different ISIs was similar to the cutaneous conditioning, lasting for ∼ 150 ms and peaking near 80 and 60 ms in the two participants (Figs. 3C and D). When applied alone the conditioning TMS did not produce an increase in the mean firing rate of the tonically active motor units (PSF in Figs. 4E and F) or in the rectified surface EMG (Figs. 3G and H), again suggesting that the facilitation of the H-reflex was not due to postsynaptic changes in the motoneurons.

**Figure 4.**
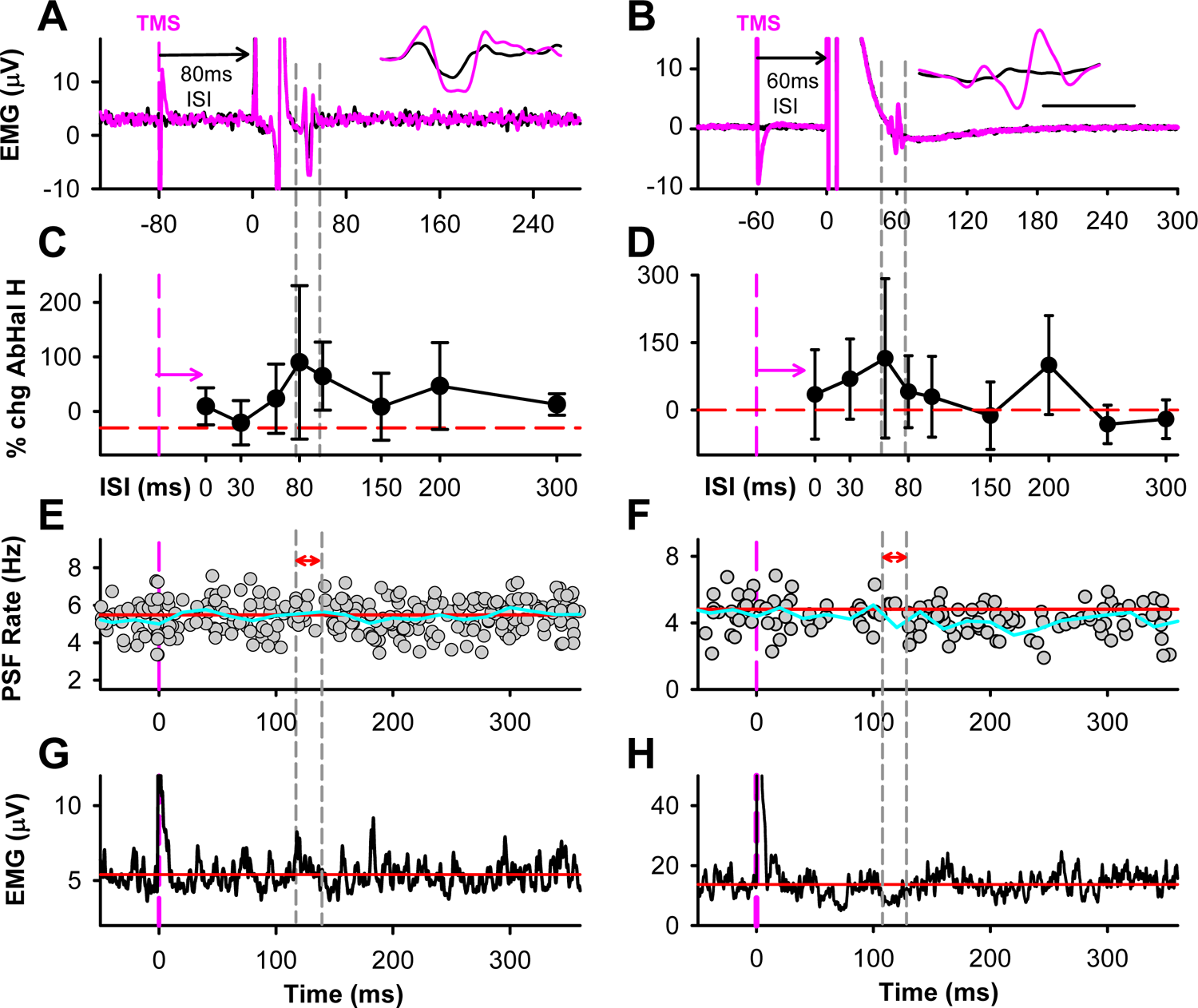
Short-duration H-reflex facilitation by CST inputs. A&B) Similar format to Figure 2 with example of test (black) and conditioned (pink) AbHal unrectified EMG illustrating conditioned H-reflex from TMS (0.9xAMT) in two participants at the 80 ms ISI in A and 60 ms ISI in B. **C&D)** Mean (± SD) % change of conditioned AbHal H-reflex at each ISI. **E&F)** PSF of a AbHal single motor unit, time-locked to TMS at 0 ms (vertical pink line) with 139 sweeps in E and 86 sweeps in F. **G&H)** Averaged rectified AbHal EMG recorded with TMS conditioning stimulation alone with 71 sweeps in G and 37 sweeps in H.

In the group average (n = 9 participants), the AbHal H-reflex was facilitated across the various ISIs with TMS conditioning (Fig. 5Ai, see statistics in legend) similar to the sensory afferent conditioning of the H-reflex, being significant at the 80 ms ISI (51.1 ± 41.8%, *p* = 0.033), and similar in duration to the expected CST-evoked phasic PAD (Carpenter *et al*., 1963). In contrast, the PSF and background EMG were not altered by the conditioning TMS (Figs. 5Aii&Aiii), indicating a lack of direct effect from the TMS on the AbHal motoneurons, similar to the low intensity CPN and cDPN stimulation. The small (but insignificant) increase in EMG at the 80 ms bin was due to motor unit synchronization from the preceding small EMG suppression and thus, not a reflection of motoneuron depolarization at this time. The maximum facilitation of the AbHal H-reflex across the various ISIs in each participant was 71.2 ± 33.7% (*p* < 0.001, Fig. 5B), with no change in the firing rate of the AbHal motor units or rectified EMG at time points when the maximal conditioned H-reflex occurred at the spinal cord. The soleus and vastus lateralis (Quad) H-reflexes were also facilitated by a low intensity TMS pulse (Fig. 5C). The peak facilitation of the soleus H-reflex was 37.6 ± 29.5% (*p* < 0.001, n = 9) and Quad H-reflex was 104.4 ± 144.5% (*p* = 0.012, n = 3) at an average ISI of 78.0 ± 24.4 ms for both muscles (not shown), with no corresponding change in PSF or EMG (*p* all > 0.50).

**Figure 5.**
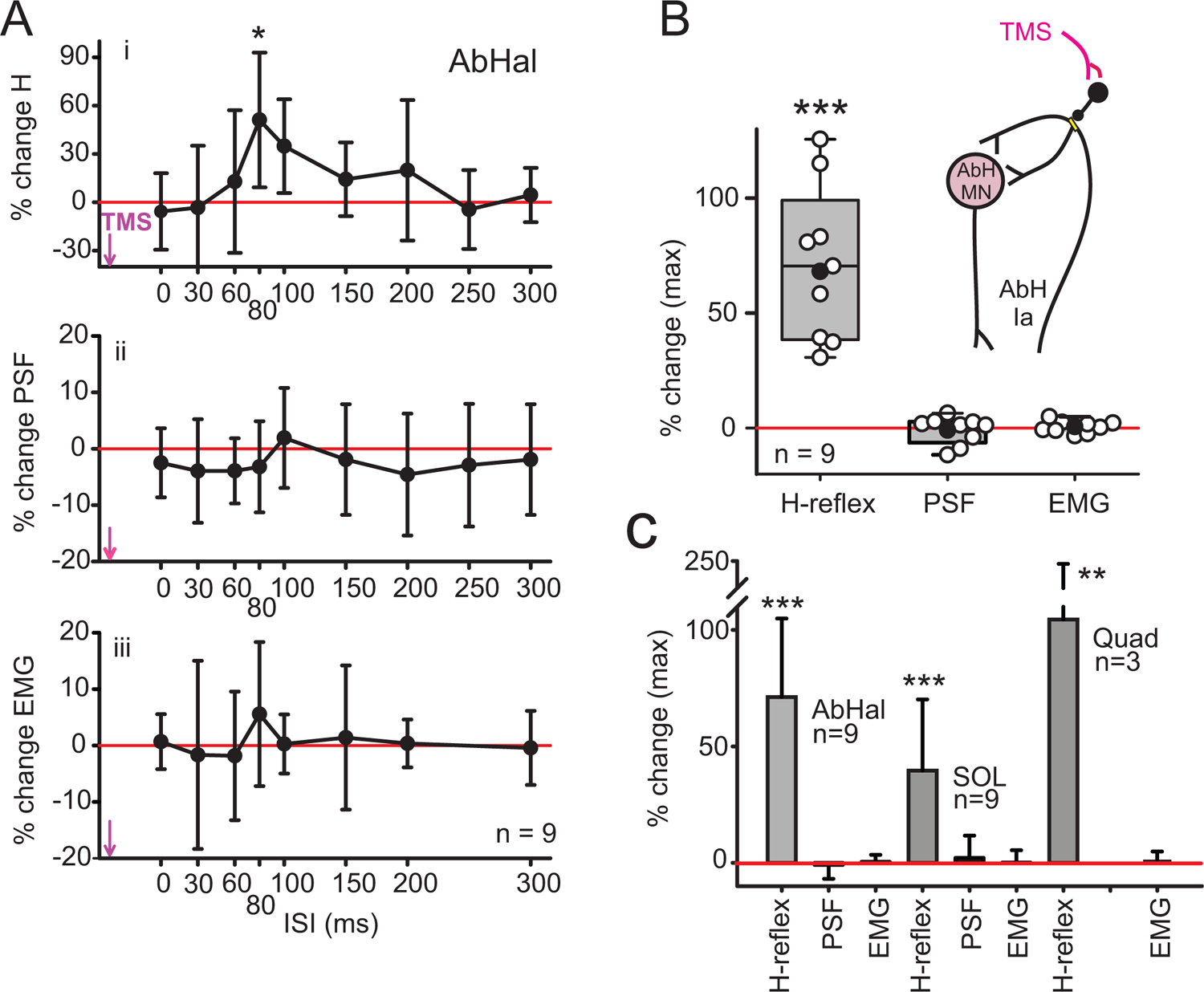
Short-duration H-reflex facilitation by CST inputs. Group data. A) Same format to Figure 3. Mean % change in (i) AbHal H-reflex (ii), motor unit firing rate (PSF) and (iii) rectified EMG from a prior TMS conditioning (0.9xAMT) at each ISI across the group (n=9). The average size of the test AbHal H-reflex was 0.11±0.82 mV (Mmax was not recorded in this muscle). Overall there was an effect of the condition-test ISI on the AbHal H-reflex [F(8,9) = 3.345, *p* = 0.002, one-way ANOVA] but not for the PSF [F(8,7) = 0.630, *p* = 0.73, one-way ANOVA] or EMG [F(8,7) = 0.501, p < 0.83), one-way ANOVA]. **B)** Left bar: Peak % change of the conditioned AbHal H-reflex. Middle bar: % change PSF at the ISI time bin where the peak conditioned H-reflex would have been activated at the spinal cord (all units were recorded using surface EMG). Right bar: % change of rectified surface EMG as in PSF. The mean PSF (−0.89 + 6.0%, *p* = 0.20) or EMG (0.44 ± 2.7%, *p* = 0.69) was not different from a 0 %change. **C)** Maximal % change in TMS conditioned AbHal, soleus (SOL) and vastus lateralis (Quad) H-reflexes and corresponding % change in PSF and EMG (PSF not available for Quad). Mann-Whitney U test used for pairwise comparisons to 0 %change with * p < 0.05, ** p < 0.01, *** p<0.001 representing statistically significant difference.

### Cutaneous and proprioceptive facilitation of H-reflexes to produce tonic PAD

*Slow (0.2 Hz) vs moderate (2 Hz) speed of cutaneous afferent trains:* We next examined if a more intense and longer duration of cutaneous DPN (cDPN) stimulation could produce a longer-lasting facilitation of the H-reflex indicative of the tonic PAD in Ia afferents produced by such stimuli (Lucas-Osma *et al*., 2018; Hari *et al*., 2021). The cDPN intensity was increased to 2 x radiating threshold (10.6 ± 1.7mA) to potentially produce GABA spillover and applied for 10 s at either 2 Hz or 0.2 Hz to determine if there was a frequency-dependent effect, as previously shown for tonic PAD mediated by α5 GABA_A_ receptors (Lucas-Osma *et al*., 2018). During both 10 s stimulation trains (2 or 0.2 Hz), the soleus H-reflex was facilitated (pink circles at 0 s time bin, Fig. 6A) compared to the preconditioned test H-reflexes (< 0 s time bins), with direct motoneuron depolarization occurring during the period of cDPN stimulation (not shown). The H-reflexes following the 2 Hz or 0.2 Hz conditioning cDPN stimulation trains (> 0 s time bins) increased across all binned time points (*p* < 0.001, see statistics in legend) with post-hoc analysis revealing that H-reflexes were significantly larger at all time points for the 2 Hz train only (*p* < 0.05). The facilitation of the H-reflex was greater following the 2 Hz stimulation train compared to the 0.2 Hz train at several of the binned time points (*p* < 0.05, marked by *’s in Fig. 6A).

**Figure 6.**
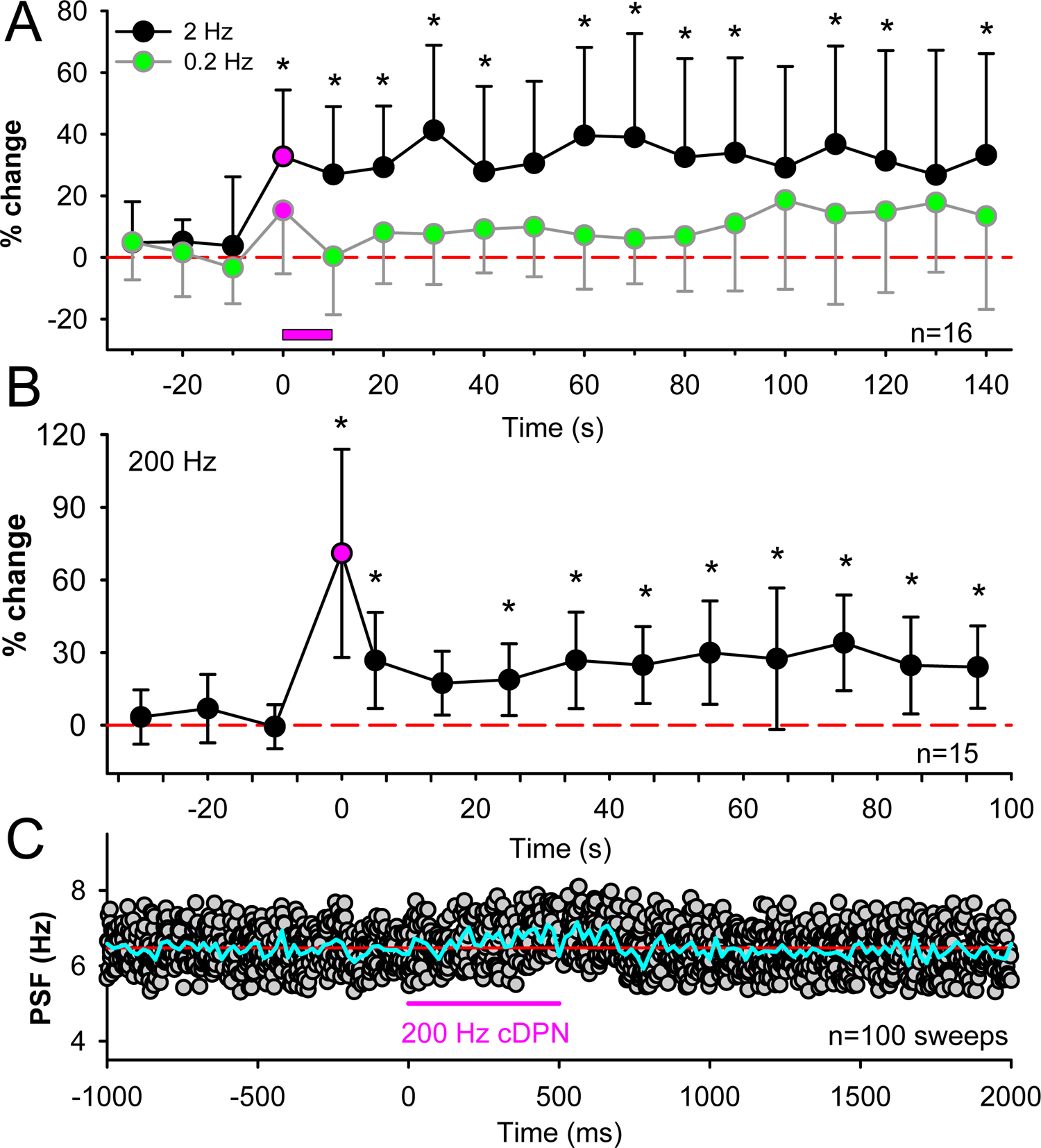
Long-duration H-reflex facilitation. A) Mean ± SD of SOL H-reflexes before (< 0 s) and after (> 0 s) a 10 s train of 2 Hz (black circles) or 0.2 Hz (green circles) cDPN stimulation shown by pink bar (n = 16). H-reflexes that were immediately preceded by a cDPN stimulation (500 ms ISI) are marked in pink at 0 ms. H reflexes were evoked every 5 seconds and data were averaged into 10-s bins. Dashed red line indicates 0 %change. H-reflexes were larger across all time points for both the 0.2 Hz (F(15,18) = 2.56, *p* <0.001) and 2 Hz (F(15,18) = 6.39, *p* <0.001) stimulation trains (one-way ANOVA) with H-reflexes after the 2 Hz stimulation greater than a 0 %change at all time bins (p < 0.05, Tukey). The increase in H-reflex was larger for the 2 Hz stimulation compared to the 0.2 Hz stimulation [F(1, 17) = 2.2, *p* = 0.005, two-way ANOVA] with * indicating when 2 Hz values > 0.2 Hz values (*p* < 0.05, Tukey). **B**) Same format in A but in response to a fast cDPN train (200 Hz for 500 ms, n=15). Pink circle represents conditioned H-reflex where cDPN stimulation preceded the H-reflex by 700 ms. There was an effect of time on the H-reflex [Chi-square = 100.6, DF = 14 (*p* <0.001)] with * indicating significant difference from a 0 %change (*p* < 0.05, Tukey). **C)** PSF of soleus motor unit in response to fast cDPN stimulation used in B. Overlay of 100 stimulation trials from one participant. Blue line is binned average of frequency points (PSF 20 ms bin width). Red line is pre-stimulus rate. `#`

*Fast (200 Hz) cutaneous afferent train:* Given the larger sustained facilitation of H-reflexes by the faster conditioning stimulation train of 2 Hz, we also tested a much higher frequency conditioning stimulation train of the cDPN at 200 Hz, but of shorter duration (500 ms). This was very effective in facilitating the H-reflex, consistent with the long-lasting tonic PAD and facilitation of monosynaptic reflexes evoked by this stimulation in animals, lasting for at least one minute after the train was terminated (Lucas-Osma *et al*., 2018; Hari *et al*., 2021). Because high frequency stimuli can be painful, we used an intensity of cDPN stimulation that was below radiating threshold (3.6 ± 0.3mA). The 500 ms, 200 Hz cDPN stimulation produced a large facilitation of the H-reflex immediately after the stimulus train (pink circle at 0 s time bin, Fig. 5B) compared to the unconditioned H-reflexes (black circles, at time bins < 0 s). This large H-reflex facilitation was associated with a ∼ 1 s increase in the PSF of the soleus motor units following the cDPN train (Fig. 6C) and thus, was partly produced by direct facilitation of the soleus motoneurons. However, after the PSF returned to baseline by 1 s, the H-reflex continued to be facilitated for at least 95 s after the fast frequency train ended. The H-reflexes were facilitated at many time points following the conditioning cDPN train (at time bins > 0 ms, marked by *, *p* < 0.05), in contrast to any of the H-reflexes evoked before the cDPN train (at time bins < 0 s).

*Slow (0.2 Hz) proprioceptive afferent train:* The small gradual increase in H-reflexes from the 0.2 Hz cDPN stimulation was likely produced by a small tonic PAD from cutaneous afferent pathways with a buildup in extra-synaptic GABA (Lucas-Osma *et al*., 2018; Hari *et al*., 2021). Similarly, a tonic PAD may also have been produced by the repeated activation of the soleus Ia afferents themselves. During repetitive stimulation of Ia afferents in the rodent, tonic PAD can be produced in other Ia afferents via the classic tri-synaptic pathway, and also in the stimulated Ia afferent itself, the latter termed self-facilitation (see Extended Fig. 10 in Hari *et al.,* 2021). Thus, we predicted that repeated activation of Ia afferents by TN stimulation alone may also produce long-lasting facilitation of the soleus H-reflex via self-facilitation from tonic PAD (see schematic in Fig. 7). To examine this, we measured if there was any buildup of *test* H-reflexes during baseline measures before any conditioning stimulation was applied. The repetition rate of every 5 seconds of TN stimulation likely produces a small amount of post-activation depression (Hultborn *et al*., 1996), but the self-facilitation of Ia afferents from tonic PAD may override this. In the 19 participants tested, H-reflexes evoked from the 1^st^ to the 14^th^ TN stimulation gradually increased in amplitude over a period of 1 minute compared to the test H-reflexes that reached a steady state after the first run of conditioning stimuli (Fig. 7, see statistics in legend, *p* < 0.001). Post-hoc, the 6^th^ to 14^th^ H-reflexes were all larger than the 1^st^ H-reflex (*p* < 0.001), suggesting self-facilitation of the Ia afferents arising from the repeated TN stimulation.

**Figure 7.**
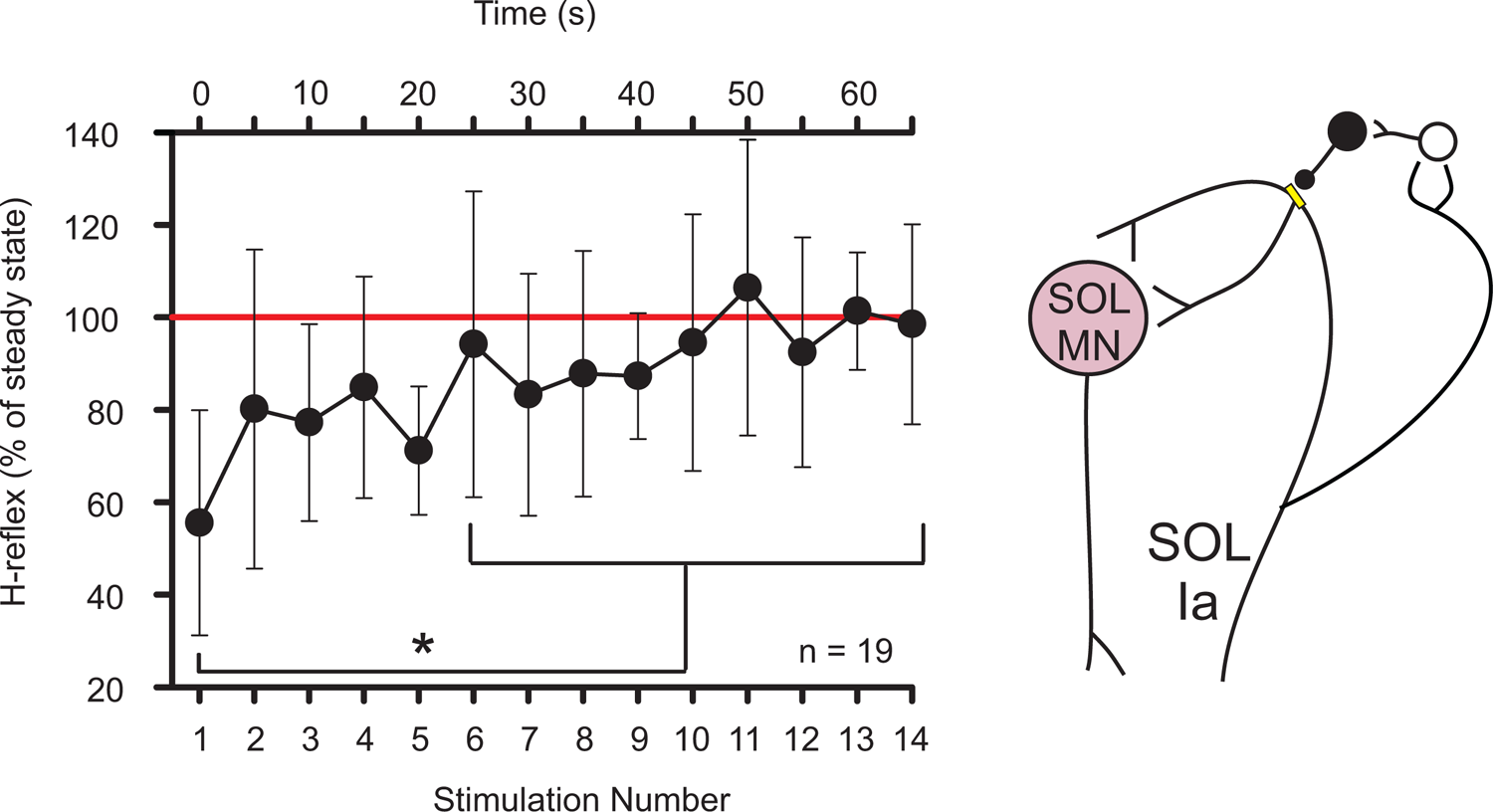
Self-facilitation of H-reflex during tonic PAD. Left: Mean ± SD of first 14 test SOL H-reflexes before any conditioning stimulation was applied averaged across 19 participants. H-reflexes were evoked every 5 seconds at ∼ 30% of maximum H-reflex. H-reflexes were expressed as a % of the average of the 7 H-reflexes following the first run of conditioned H-reflexes in set 2 (see Fig. 2, set 2 average = 100% as marked by red horizontal line). The test H-reflexes increased over the 14 TN stimulations (F(18,13) = 3.93, *p* < 0.001, one-way ANOVA) with H-reflexes 6 to 14 larger than the first H-reflex (*p* < 0.001, Tukey). Right: schematic of SOL Ia afferent collateral activating PAD circuit that synapses back onto its own branchpoint node to produce tonic PAD and self-facilitation.

### Cutaneous facilitation of single motor unit discharge probability

To provide stronger evidence that a conditioning cutaneous stimulus facilitates the soleus H-reflex by preventing intermittent failure in Ia afferent conduction and generation of the EPSP on the motoneuron, we examined whether the firing *probability* of soleus motor units within the H-reflex (Ia-EPSP) window was increased without producing an increase in the *amplitude* of the motoneuron EPSP estimated by the PSF method. The PSF requires many motor unit firing trials to be averaged and thus we used cutaneous stimulation trains that produce long-duration increases in afferent depolarization (tonic PAD), where we repeatedly tested the H-reflex before and then after conditioning. The intensity of the TN stimulation used to evoke the H-reflex was adjusted to produce a firing probability of the motor units near or below 50%, since intermittent failure in afferent branch points are more readily seen at low stimulation intensities (Hari *et al*., 2021). Participants held a weak plantarflexion to recruit a motor unit with a steady background firing rate, upon which we could evaluate changes in firing rate with TN stimulation (PSF, Fig. 8A). At the onset of the H-reflex window (marked by vertical grey line, representative data from a single participant in Fig 8A), the PSF increased with a profile consistent with the underlying Ia EPSP that produces the H-reflex (PSF in light blue), as previously detailed (Turker & Powers, 2005). A brief duration (50 ms), low intensity cDPN stimulation train (4.0 ± 0.55 mA, 200 Hz) was applied 500 ms before the TN stimulation to avoid any postsynaptic effects on the motoneuron at the time of Ia activation and allow motor units to be reliably followed. By ∼200 ms after the conditioning cDPN train, any depolarization of the SOL motoneuron subsided as reflected in the PSF returning to baseline before the H-reflex was evoked (light blue trace near red line, Fig. 8Aii). In this participant, the probability of motor unit discharge within the H-reflex window (15 ms in duration) increased from 58% during the test-alone trials (Fig. 8Ai) to 71% during the cDPN-conditioning trials (Fig. 8Aii). The increased probability of motor unit discharge occurred even though the PSF, representing the profile of the Ia-evoked EPSP (∼ 15 ms in duration in rats, Hari et al., 2021), was not altered by the cutaneous conditioning stimulation (note overlay of blue test PSF and pink conditioned PSF, inset of Fig. 8Aii).

**Figure 8.**
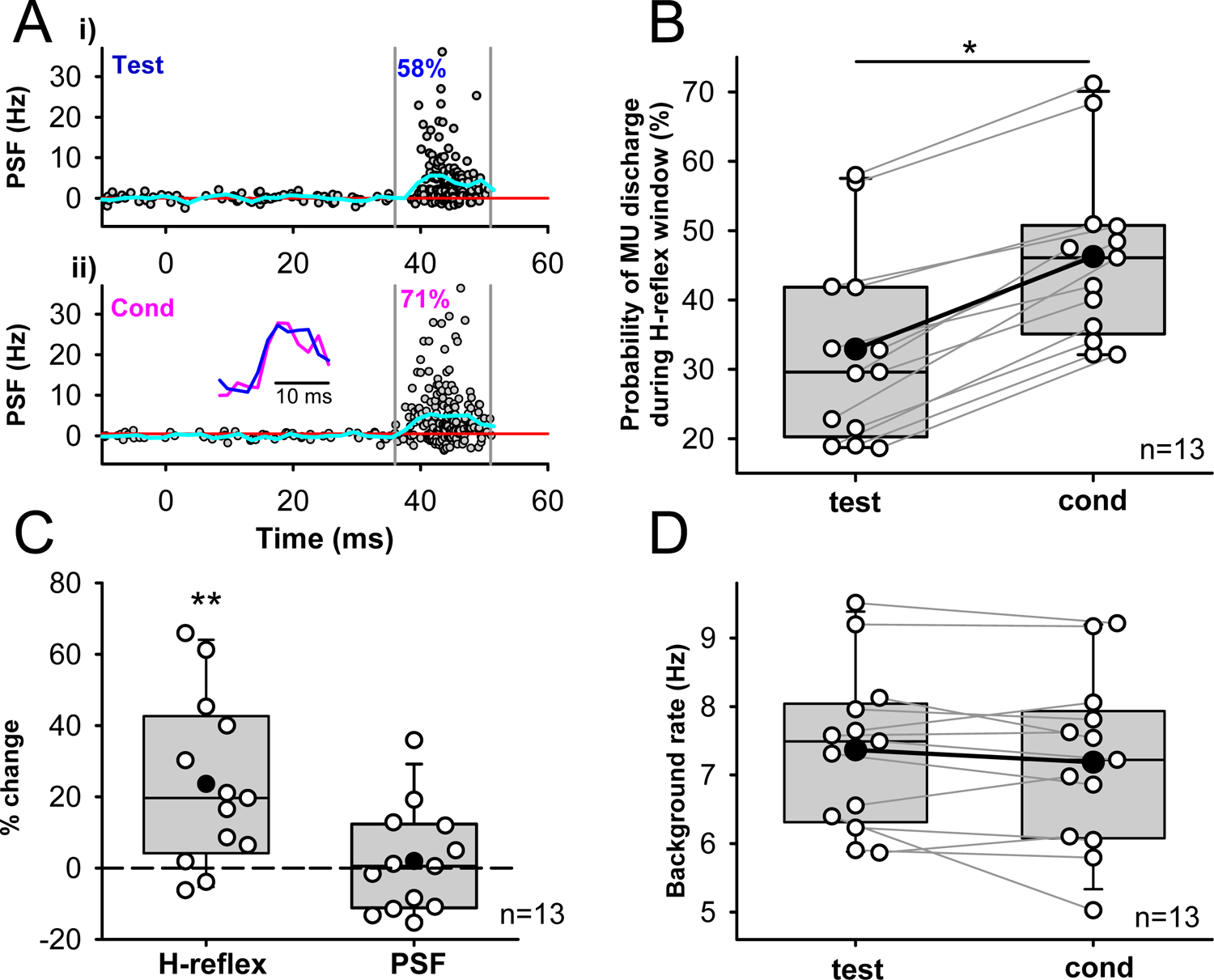
Probability of Single Motor Unit Discharge. A) A representative PSF of SOL motor units recorded from a single male participant using HDsEMG, time-locked to the TN stimulation (at 0 ms) without (**i**, test) and with (**ii**, cond) cDPN conditioning (10 pulses, 200 Hz, 500 ms ISI). Activity of 11 units are superimposed over 302 sweeps in i (test H-reflexes) and 311 sweeps in ii (conditioned H-reflexes). To reduce variability across units, the mean pre-stimulus firing rate was subtracted from each unit. Mean PSF (blue line) is plotted over mean pre-stimulus rate (red line). The firing probability of the units were measured within the H-reflex window between the grey vertical lines. *Inset:* estimated EPSP from the test (blue) and conditioned (pink) PSFs. **B)** Probability of motor unit (MU) discharge during H-reflex window before (test) and after (cond) cDPN conditioning for each participant (white circles, n = 13), mean represented by the black circle and median by the horizontal line, 25th and 75th percentiles by the box bounds, and the 95th and 5th percentiles by whiskers. Unit activity was measured with HDsEMG in 12 participants and with intramuscular EMG in 1 participant. Mann Whitney U test was used for pairwise comparison between test and conditioned trials with * representing statistically significant difference (*p* < 0.05). **C)** % change of the conditioned SOL H-reflex (left bar) and % change of the PSF within the H-reflex window (right bar), with the H-reflex greater than a 0 % change (23.6 ± 23.6%, *p* = 0.001 noted by **) but no change in PSF (1.9 ± 14.9% change, *p* = 0.74), both using Mann Whitney U test. **D)** Average background firing rate of the SOL motor units measured 100 ms before TN nerve stimulation from the test (left bar) and conditioned (right bar) stimulation trials, with no difference between the two (*p* = 0.71, Student’s t-test).

Across the group (n = 13), the firing probability of the motor units within the test H-reflex window was 32.9 ± 14.2 % and increased to 46.3 ± 12.8% during the cutaneous conditioning trials (*p* = 0.014, Fig. 8B). Importantly, the average firing rate of the PSF within the H-reflex window (13.6 ± 4.6 ms in duration; reflecting the EPSP size) did not change in response to the cutaneous conditioning (Fig. 8C right bar), suggesting that an increase in firing probability of the soleus motoneurons activated by the Ia afferents occurred without an increase in the amplitude of the EPSP. In 8 of the 13 participants, the mean PSF was unchanged or even decreased in response to the cutaneous conditioning (Fig. 8D), ruling out changes in motoneuron facilitation (postsynaptic) accounting for the increased motor unit firing probability. The overall soleus H-reflex measured from the surface EMG also increased with conditioning (Fig. 8C left bar), as expected from Figure 6, consistent with the increased probability of the EPSP dominating over any changes in EPSP size and leading to a net increase in the H-reflex. Across the group, the mean background firing rate of the motor unit PSF 100 ms before the TN stimulation with cutaneous conditioning (cond, right bar Fig. 8D) was not different compared to without conditioning (test, left bar) in all participants, further supporting the conclusion that changes in firing probability of the units were not mediated by postsynaptic facilitation of the soleus motoneurons.

## Discussion

Over the past 60 years, GABA and the depolarization it produces in peripheral afferents (PAD), was thought to inhibit sensory transmission in the spinal cord by reducing action potential size at the afferent terminal and subsequent neurotransmitter release. However, in the brainstem (Calix of Held), cerebellum (Purkinje cells) and cortex (basket cells) where it is sometimes possible to record directly from presynaptic boutons, the depolarization of the axon terminal from GABA_A_ or glycine receptors can facilitate neurotransmitter release (typically by increasing intracellular calcium) and subsequent excitatory postsynaptic currents [reviewed in (Trigo *et al*., 2008; Zorrilla de San Martin *et al*., 2017)]. We do not think that this terminal facilitation accounts for the facilitation of the monosynaptic reflex detailed here or in the related animal studies (Hari *et al*., 2021), because GABA_A_ receptors are mostly absent from proprioceptive afferent terminals in this reflex pathway. Instead, facilitation of Ia mediated EPSPs in the motoneuron likely occurs when axon nodes are depolarized from the activation of nodal GABA_A_ receptors, which helps to bring sodium channels closer and more rapidly to threshold to reduce branch point failure in the Ia afferent (Hari *et al*., 2021). This depolarization, or PAD, measured in rodent Ia afferents is readily activated by other Ia afferents or cutaneous and pain afferents demonstrating a facilitatory, rather than inhibitory, role of sensory inputs on the conduction of action potentials in proprioceptive axons. In agreement with this animal study, we provide evidence that the conduction in Ia afferents is similarly facilitated by proprioceptive, cutaneous and CST inputs in the human.

### Short-duration facilitation of H-reflexes by sensory and CST pathways

The profile of H-reflex facilitation from a brief conditioning stimulation of sensory or CST pathways lasted for about 100-200 ms and peaked near the 60 or 80 ms ISI. A similar profile of monosynaptic reflex facilitation was produced in rodents following either a brief cutaneous stimulation or from direct light activation of GABA_axo_ interneurons (Hari *et al*., 2021). In both cases, we propose that the time course of monosynaptic reflex facilitation follows the time course of the evoked phasic PAD in the Ia afferent. An illustration of the general profile of H-reflex facilitation in relation to the estimated PAD evoked in the Ia afferents is provided in Figure 1. For instance, at the 0 ms ISI, the phasic PAD from the conditioning stimulation was not yet activated in the Ia afferents when the afferent was activated by the TN stimulation for the H-reflex. This likely produces a motoneuron EPSP and H-reflex that is uninfluenced by PAD. However, when the TN stimulation followed the conditioning stimulation by 60 to 80 ms, the activation of the TN Ia afferents occurs during the presence of the PAD, allowing the TN stimulation to activate more Ia afferent branches and produce more or larger EPSPs and a larger H-reflex (H-reflex at 60 to 80 ms ISI > at 0 ms ISI). An axonal (nodal) mechanism of H-reflex facilitation is likely because when the conditioning stimulation was applied alone, the PSF was slightly below the mean pre-stimulus rate, indicating that the conditioning stimulation itself did not depolarize the motoneurons to facilitate the H-reflex.

Although the mean profile of H-reflex facilitation closely follows the estimated profile of phasic PAD from the 60 or 80 ms ISI and onwards, the facilitation of the H-reflex at the earlier ISIs are smaller than expected based on the PAD profile. This may be due to direct effects on the motoneurons from the conditioning stimulation that mask the facilitation of Ia transmission by PAD. For example, any excitatory or inhibitory activation of the motoneuron may have prevented full H-reflex facilitation at these earlier ISIs due to postsynaptic shunting or direct inhibition of the motoneuron as shown in rodents (Hari *et al*., 2021). Small decreases in the PSF during these earlier ISIs provide some evidence that the motoneuron may have been slightly inhibited by the conditioning sensory and CST stimulation. In the rodent, when the direct effects on the motoneuron from the conditioning stimulation are removed with voltage clamp, the full effect of the PAD facilitation on monosynaptic reflexes is unmasked. Thus, if anything subtle postsynaptic effects on the motoneuron from the conditioning stimulation tend to decrease the Ia EPSP, strengthening the conclusion that any H-reflex facilitation occurred from increases in Ia afferent conduction.

Both cutaneous and CST inputs have the majority of their terminations in the dorsal horn as measured from anatomical tracings (Lucas-Osma *et al*., 2018; Ueno *et al*., 2018). These inputs may activate the tri-synaptic GABA pathway, possibly including activating the first order glutamatergic interneurons that synapse onto the GABA_axo_ interneurons in this pathway, which in turn activate GABA_A_ receptors on dorsal nodes of the Ia afferent (Hari *et al*., 2021). Similar to pain afferents (Hayes & Carlton, 1992), terminals of the CST also synapse directly onto the GABA_axo_ interneurons (identified as d14: GAD2+/Ptf1a+ neurons) that project onto Ia afferents (Ueno *et al*., 2018). As a descending regulator of sensory inflow to the brain and spinal cord (Liu *et al*., 2018), our results indicate that CST projections that activate GABA_axo_ interneurons directly facilitate Ia afferent conduction and reflex activation of motoneurons during movement.

The net facilitating action of GABA on afferent conduction is likely due to the relatively greater expression of GABA_A_ receptors on the dorsally located nodes of the myelinated segments of Ia afferents, compared to the sparser receptor expression found on the unmyelinated terminals of these Ia afferents (Lucas-Osma *et al*., 2018; Hari *et al*., 2021). The few GABA_A_ receptors at the terminals could, in principle, provide a graded shunting of current to produce presynaptic inhibition of Ia inputs onto the motoneuron. However, mathematical models demonstrate that this shunting is not sufficient to reduce the size of the action potential invading the terminal (Walmsley *et al*., 1995; Hari *et al*., 2021). Moreover, PAD at the terminal is small (Lucas-Osma *et al*., 2018), likely owing to the small number of terminal GABA_A_ receptors (Alvarez *et al*., 1996; Betley *et al*., 2009; Fink *et al*., 2014) compared to dorsal parts of the afferent (Lucas-Osma *et al*., 2018) and the large electrotonic attenuation of current from the last node to the terminal (Hari *et al*., 2021). However, GABA may also activate GABA_B_ receptors on the afferent terminal and GABA_A_ receptors on the motoneuron to reduce the size of the monosynaptic reflex (Pierce & Mendell, 1993; Hughes *et al*., 2005; Hari *et al*., 2021). Thus, the net increase in monosynaptic reflexes from the activation of GABA_axo_ interneurons is likely mediated by the activation of GABA_A_ receptors on the dorsal regions of the Ia afferent that have a stronger facilitatory effect on Ia afferent conduction compared to the inhibitory effect of GABA_B_ receptors activated on afferent terminals and the GABA_A_ receptors on the motoneurons. This balance may favor the facilitation of reflexes when the conditioning stimuli are moderate or small and instead favor a suppression of H-reflexes when higher intensity conditioning stimuli are applied, which may have stronger effects on GABA_B_ receptor-mediated presynaptic inhibition and/or direct motoneuron inhibition (Hari *et al*., 2021). It remains to be determined if the same or separate GABA_axo_ neurons innervate nodes (GABA_A_) and terminals (GABA_B_). For example, perhaps a separate group of GABAergic neurons innervate the terminals (and GABA_B_ receptors) and are driven by homonymous nerve stimulation, causing the strong depression of the H-reflex with repeated stimulation (i.e., rate dependent or post-activation depression), and another more dorsal group of GABAergic neurons mediates nodal facilitation (via GABA_A_ receptors) that are driven by more diverse afferent (e.g., cutaneous) and descending inputs.

### Long-lasting facilitation of Ia afferents by cutaneous and proprioceptive inputs

Our results demonstrate that trains of cutaneous stimulation (0.2 Hz to 200 Hz) facilitate the soleus H-reflex for up to 2 minutes, similar to the duration of long-lasting (tonic) PAD recorded in rodent Ia afferents in response to identical stimulation trains applied to a dorsal root (Lucas-Osma *et al*., 2018; Hari *et al*., 2021). The long duration of PAD evoked in Ia afferents from the multiple, especially high frequency, sensory inputs is produced by the activation of extra-synaptic 5 GABA receptors on the Ia afferent nodes, potentially from GABA spillover produced by the repeated activation of GABA_axo_ interneurons (Lucas-Osma *et al*., 2018). It is unlikely that the motoneuron is continually facilitated for 2 minutes by these high frequency, 10 second stimulation trains given that the membrane potential of the motoneuron in the rat, and the motor unit firing rates in the human (PSF), return to pre-stimulation baseline by less than 1 second after the stimulation train. The trains of low frequency (0.2 Hz) cutaneous stimulation produce a smaller sustained facilitation of the H-reflex, comparable to the low-amplitude tonic PAD produced from the same stimulation train in the rodent (Lucas-Osma *et al*., 2018), and thus this small facilitation is likely due to a small tonic PAD. It is likewise possible that the gradual increase in H-reflexes over the 2-minute recording is produced by tonic PAD evoked by the repeated (0.2 Hz) activation of the soleus Ia afferents themselves when evoking the H-reflex (repetitive TN stimulation). The gradual increase in test H-reflexes with TN stimulation alone supports this hypothesis of self-facilitation where collaterals of the Ia afferent activate a PAD network that synapses back onto its own branch point nodes. Self-facilitation of the Ia afferents is most readily revealed when we use low intensities of TN stimulation that produced an H-reflex of ∼30% of maximum, giving headroom for recruiting new afferent branches with tonic PAD (Hari *et al*., 2021).

### Probability of motor unit firing

To strengthen the conclusion that the facilitation of the H-reflex by the sensory or CST conditioning is mediated by facilitation of the Ia afferents, we found it useful to measure single motor unit (motoneuron) activity before and during the H-reflex (Ia-EPSP) window. This allows us to: 1) examine whether the conditioning stimulation alone changes the motoneuron depolarization by examining baseline motor unit firing rates just before the H-reflex (postsynaptic actions), 2) examine whether the conditioning changes the EPSP size in a graded manner by measuring the PSF during the Ia-EPSP (H-reflex) window since graded changes in EPSP would be mediated by changes in either presynaptic inhibition or postsynaptic facilitation and 3) examine the probability of the evoked Ia-EPSP (all-or-nothing failure) reflected in whether the motor unit participated in the H-reflex or not. Overall, we found that the conditioning-evoked PAD was not associated with an increase in baseline motor unit firing or the size of the estimated EPSP, consistent with a lack of postsynaptic facilitation or decrease in presynaptic inhibition that would otherwise grade the EPSP size. If anything, conditioning tended to slightly slow motor unit firing or hyperpolarize motoneurons, as in rats (Hari *et al*., 2021), which would decrease the probability of the motor unit contributing to the H-reflex. In contrast, we found that the conditioning consistently increased the probability of the motor unit participating in the H-reflex, in agreement with conclusions from rats that PAD prevents branch point failure and reduces the probability of intermittent, all-or-nothing EPSP failures.

Low amplitude TN stimulation intensities were used with the intention of evoking monosynaptic soleus H-reflexes only and not polysynaptic reflexes. Thus, the PSF within the 15 ms H-reflex window likely reflected motor unit discharge during the monosynaptic EPSP and its increased probability by increases in Ia afferent conduction. In support of this, mechanical cutaneous conditioning of the H-reflex in a forearm muscle also increased the discharge probability of motor units measured during the first 0.5 ms of the reflex window (i.e., during the early monosynaptic component of the EPSP), without increases in integrated EMG and motor unit firing rates, again showing that the motoneuron was not facilitated by cutaneous conditioning but the firing probability of the Ia afferents were (Aimonetti *et al*., 2000). In addition, the Aimonetti *et al.,* 2000 results were obtained using single motor unit recordings from intramuscular EMG and corroborates our findings using decomposition of single motor units from HDsEMG during reflex activity (see also Yavuz *et al*., 2015).

Outside of sensory-evoked PAD preventing branch point failure, there are a few other possible explanations for the increased probability of motor units contributing to the H-reflex during conditioning which we must rule out. The conditioning input might somehow increase the probability of quantal transmitter release at the terminal, thereby increasing the probability of the EPSP. This could occur by a yet undescribed non-GABAergic innervation of the Ia afferent terminal that may facilitate intracellular calcium and neurotransmitter release. GABAergic effects on the afferent terminal seem unlikely to increase firing probability because imaging in rodents indicate that Ia afferent terminals mainly only express GABA_B_ receptors (Hari *et al*., 2021) which decrease, rather than increase, transmitter release via its inhibitory Gi protein coupled pathways (Curtis & Lacey, 1998). Even if a few GABA_A_ receptors were on the terminals, these have little practical depolarizing action as shown from direct terminal recordings (Lucas-Osma *et al*., 2018) and, if anything, likely inhibits transmitter release (although see Trigo *et al*., 2008). Thus, we conclude that the most likely explanation for the increased H-reflex and motor unit probability during conditioning is a PAD-mediated facilitation of branch point conduction in the afferents mediating the H-reflex. Further work is needed to confirm this, including showing that when the motor unit fires during the H-reflex window, there is a uniformly bigger H-reflex than when the motor unit fails to fire during the reflex, consistent with recruitment of unitary EPSPs that result from an afferent branch that is recruited into action.

### Relation to previous animal and human studies

The idea that cutaneous and CST pathways facilitate H-reflexes by increasing Ia afferent excitability from PAD is not at odds with previous cat data (Rudomin *et al*., 1983), but is at odds with the previous conclusion that these pathways reduced PAD measured in extensor afferents (Rudomin *et al*., 1983). In the cat studies, the size of PAD was indirectly measured by quantifying the threshold current needed to maintain a set level of antidromic firing probability of the Ia afferent when activated by extracellular stimulation in the intermediate or ventral motor nucleus in the spinal cord (Wall, 1958; Carpenter *et al*., 1963; Willis *et al*., 1976; Rudomin *et al*., 1983). Cutaneous and CST inputs reversed the lowering of the threshold current produced by tonic flexor afferent stimulation, leading to the conclusion that cutaneous and CST inputs reduced PAD and hence, the assumed presynaptic inhibition. However, this indirect measure of inhibition of PAD may have been in error, since it is in contrast to the large PAD directly observed following cutaneous afferent stimulation measured intra-axonally in more dorsal regions of the Ia afferent (Lucas-Osma *et al*., 2018; Hari *et al*., 2021) and the dorsal root potentials evoked from low threshold stimulation of the dorsal cutaneous nerve in mice (Zimmerman *et al*., 2019). To reconcile these differences, it may be that cutaneous and CST inputs activate GABA_axo_ interneurons to produce PAD in dorsal portions of the Ia afferent but at the same time, inhibit GABA_axo_ interneurons with connections to more ventral portions of the afferent (i.e., again two separated populations of interneurons). This is supported by the finding that antidromic potentials activated by stimulation of afferents in the dorsal horn are strongly facilitated by cortical stimulation in contrast to antidromic potentials evoked from stimulation of afferent terminals in the ventral horn (Carpenter *et al*., 1963). In this way, cutaneous pathways (and potentially CST pathways) may enhance dorsal nodes to secure (facilitate) action potential transmission and at the same time, reduce PAD at more ventral nodes to reduce any lowering of threshold currents. Direct measurements of PAD in dorsal and ventral parts of the Ia afferent and its modulation from cutaneous and CST inputs are needed to sort out this discrepancy (though see Lucas-Osma *et al*., 2018).

Based on the cat work, human studies have also proposed that cutaneous and CST inputs reduce the amount of PAD and presynaptic inhibition in agonist Ia afferents, the latter measured from the suppression of H-reflexes by antagonist afferents (Berardelli *et al*., 1987; Iles & Roberts, 1987; Nakashima *et al*., 1990; Iles, 1996; Meunier & Pierrot-Deseilligny, 1998; Aimonetti *et al*., 2000). In these studies, it has been proposed that the suppression of H-reflexes by antagonist afferents is reduced by cutaneous and CST pathways via dis-facilitation of the GABA_axo_ interneurons mediating PAD, which would then result in a decrease of presynaptic inhibition. However, recent evidence in rodents shows that the suppression of monosynaptic reflexes by afferents is not mediated by presynaptic inhibition of Ia afferents by PAD (Hari *et al*., 2021). Rather, inhibition of monosynaptic reflexes by afferent conditioning is produced by other mechanisms such as terminal GABA_B_ receptor activation, post-activation depression and/or postsynaptic shunting on the motoneuron (Curtis & Lacey, 1994; Walmsley *et al*., 1995; Trigo *et al*., 2008; Howell & Pugh, 2016; Zbili & Debanne, 2019; Hari *et al*., 2021). Thus, the reduced H-reflex inhibition from cutaneous and CST conditioning in previous human studies is likely explained by facilitating action potential propagation in the Ia afferents by reducing branch point failure to counteract the inhibition of the monosynaptic reflex from these other inhibitory mechanisms.

### Functional implications

Activation of GABAergic networks in the spinal cord can have both facilitating and inhibitory actions on afferent transmission within the spinal cord. We demonstrate here that proprioceptive, cutaneous and CST pathways have a net excitatory influence on Ia afferents and this may also apply to other afferent modalities as well, such as touch and pain (Lucas-Osma *et al*., 2018; Hari *et al*., 2021). Thus, the drive from the CST during volitional movements and the coincident activation of movement-related proprioceptive and cutaneous inputs may help to secure propagation of action potentials in sensory axons toward both the brain and spinal cord to facilitate the use of this sensory information in movement generation and control. Such regulation of afferent conduction may be affected by brain or spinal cord injury given the known changes to GABAergic networks following these insults (Faist *et al*., 1994; Tillakaratne *et al*., 2000; Kapitza *et al*., 2012; Mende *et al*., 2016; Khalki *et al*., 2018; Lalonde & Bui, 2021). Perhaps some of the problems with movement control and development of spasticity from injury may be produced by alterations in GABAergic control of nodal facilitation in afferents, a topic we are currently exploring.

## Additional Information

### Competing Interests

The authors have no competing interests to declare.

### Author contributions

K.M., Y.L., D.J.B and M.A.G. conceived and designed research; K.M., I.C-M., Y.L., D.J.B. and M.A.G. performed experiments; K.M., I.C-M. and M.A.G. analyzed data; K.M., B.A., C.T. and F.N. decomposed the HDsEMG data; K.M., Y.L., D.J. B. and M.A.G. interpreted results of experiments; K.M., I.C-M and M.A.G. prepared figures; K.M.. D.J.B. and M.A.G. drafted, edited and revised manuscript.

All authors approve the final version of the article and agree to be accountable for all aspects of the work. The authors confirm that all persons designated as authors are qualified.

## Funding

This work was supported by a National Science and Engineering Grant 05205 to M.A.G. and studentship funding to K.M. from the Neuroscience and Mental Health Institute and Faculty of Medicine and Dentistry at the University of Alberta.

## Acknowledgments

We thank Ms. Jennifer Duchcherer for technical assistance.

## Notes

### Competing Interest Statement

The authors have declared no competing interest.

